# Identification of Potential Hub Genes and Therapeutic Targets in Colorectal Cancer Using Integrated Bioinformatics Approaches

**DOI:** 10.1101/2024.11.16.623927

**Authors:** Kamalakannan D, Manivannan R, Suresh Gopal Kumar, Dilip Kumar

**Affiliations:** EXCEL COLLEGEOF PHARMACY,KOMARAPALAYAM; EXCEL COLLEGE OF PHARMACY,KOMARAPALAYAM

**Keywords:** Colorectal cancer (CRC), Hub genes, Bioinformatics, Prognostic markers, differentially expressed genes (DEGs)

## Abstract

In this study, we took a comprehensive approach using bioinformatics to uncover potential therapeutic targets for colorectal cancer (CRC). We started by analyzing gene expression data from GEO2R to identify genes that were differentially expressed in CRC. Then, using FunRich software, we created Venn diagrams to visualize these genes. From the 191 upregulated genes we found, we focused on potential “hub genes” by looking at their network connections and strength, using the STRING database.To understand the roles of these hub genes, we performed functional analyses like Gene Ontology (GO) and pathway enrichment through the DAVID platform. This helped us pinpoint key biological processes and pathways linked to the genes we identified. We also looked at patient survival data from GEPIA, along with information on gene expression related to disease stages and metastatic progression. This helped us identify which hub genes were most relevant for CRC.In addition, we examined genetic changes and gene expression patterns in CRC patients through databases like cBioPortal and the Human Protein Atlas. This gave us more evidence supporting the involvement of these genes in the disease. Ultimately, our analysis highlighted CXCL8, FOXC1, ICOS, and MCF2 as potential hub genes with important roles in CRC. These genes may serve as useful biomarkers for both diagnosing CRC and predicting patient outcomes, and they could also help guide the development of targeted treatments to improve survival rates.

## Introduction

Colorectal Cancer (CRC) represents a major global health issue, ranking as the third most common cancer and the second leading cause of cancer related mortality, with approximately 1.9 million new cases and 0.9 million deaths reported in 2020In India, colorectal cancer is more commonly found in men than in women. Studies show a higher incidence of signet ring carcinoma (13%) and advanced-stage disease, including metastases affecting the liver and lungs, among patients (1)(2). Several studies show that dietary habits, such as high consumption of red meat and eggs, increase the incidence of colorectal cancer (CRC)(3). Other lifestyle factors, including low physical activity, smoking, alcohol consumption, and metabolic disorders like high body mass index, type 2 diabetes, and hypertension, have also been linked to a higher risk of CRC (4,5). The primary pathophysiology of colorectal cancer (CRC) involves chromosomal instability, microsatellite instability, and epigenetic modifications, including DNA methylation, histone modifications, and non-coding RNA changes (6) Mutations in the APC, KRAS, TP53, and MYC genes are primarily associated with tumor development in both sporadic CRC and inflammatory bowel disease-associated CRC (7).

Microarray technology, coupled with bioinformatics, enables the analysis of large datasets generated from genomics, transcriptomics, and proteomics experiments ((8,9) Advanced platforms like the Affymetrix GeneChip and Illumina BeadArray are widely used in microarray studies (10) The integration of deep learning and neural graph networks has been instrumental in predicting protein function, aiding drug discovery, and improving cancer prognosis and diagnosis (11).The NCBI Gene Expression Omnibus (GEO) is a public repository for gene expression and epigenomics data, containing over 200,000 studies and 6.5 million samples (12). It provides web-based tools for the analysis and visualization of differential expression, including new interactive graphical plots available in GEO2R (13).

The Cancer Genome Atlas (TCGA) is a comprehensive public repository of cancer genomic data, covering 33 cancer types, including rare cancers (14). It contains information on DNA sequences, transcriptional data, and epigenetic modifications, enabling the identification of genes, pathways, and accurate cancer classification (15). The TCGA dataset has supported the development of several databases, such as UALCAN (16), cBioPortal(17), STRING(18), and the Human Protein Atlas(19), for data collection and analysis. mRNA expression, networking, and DNA methylation were visualized using software tools like Cytoscape(20,21), Fun Rich(22) to identify significant prognostic markers across various cancers. This comprehensive analysis aids in recognizing tumor-related genes, and the application of microarray technology allows for an in-depth investigation into these key genes, with the aim of identifying potential molecular targets and diagnostic indicators.

Our study aimed to use bioinformatic tools to uncover key genes involved in the development of colorectal cancer (CRC) and assess their potential as treatment targets. We began by examining microarray data, identifying significantly upregulated genes associated with cancer-related pathways, particularly those linked to CRC. To get a clearer picture of CRC progression, we analyzed these genes through protein-protein interaction (PPI) networks, Gene Ontology (GO) functional classifications, and KEGG pathway mapping.

Additionally, we conducted survival and expression studies on these critical genes, focusing on how their mRNA levels varied across cancer stages, nodal metastasis, and patient survival rates. By linking survival data with gene expression patterns, we pinpointed specific hub genes that show strong potential as therapeutic targets or biomarkers for CRC diagnosis

## Materials and Methods

### Data collection

The GSE164191 dataset includes blood samples from 59 patients with colorectal cancer and 62 healthy individuals, all analyzed using the Affymetrix Human Genome U133 Plus 2.0 Array platform. This dataset, submitted by Sun Z et al (23) and available on the GEO database, allows researchers to examine gene expression in blood to spot immune-related genes specifically associated with colorectal cancer. These findings could be helpful for developing diagnostic tools that distinguish between colorectal cancer patients and healthy individuals based on blood gene expression patterns.

### Data Processing of DEGs

Differentially expressed genes (DEGs) of colorectal cancer and healthy patients were identified through GEO2R (http://www.ncbi.nlm.nih.gov/geo/geo2r), applying a statistical significance threshold of p < 0.01. Genes with a log fold change log2FC > 1 were classified as upregulated, while those with a log2FC < −1 were considered to be downregulated ((24) A volcano plot was used to visualize the DEGs, illustrating the relationship between fold change and p-value for gene expression levels.

### Gene Ontology (GO) Enrichment and Kyoto Encyclopedia of Genes and Genomes (KEGG) Pathway Enrichment Analysis

GO analysis is a widely used method for classifying genes and their RNA or protein products into specific GO categories, facilitating the identification of distinct biological characteristics from high-throughput transcriptomic or genomic data. KEGG provides a collection of databases with comprehensive information on genomes, diseases, biological pathways, drugs, and chemical substances. The Database for Annotation, Visualization, and Integrated Discovery (DAVID; http://david.ncifcrf.gov, version 6.8) is an online resource that integrates biological knowledge and analytical tools, frequently used for analyzing GO and KEGG pathways (25,26). A p-value of <0.05 was considered the threshold for significant enrichment results.

### PPI Network and CytoHubba Analysis

STRING version 10.0 (https://string-db.org/), known as the Search Tool for the Retrieval of Interacting Genes/Proteins, is a web-based resource designed to identify both physical and functional associations among genes and proteins. In this study, we used STRING to visualize the protein-protein interaction (PPI) network of differentially expressed genes (DEGs), setting a combined interaction score of greater than 0.4 as the threshold (27). Further analysis of the PPI networks was conducted with Cytoscape, where key hub genes were identified using the CytoHubba plugin, which applied 11 different algorithms. From this analysis, the top 10 most dysregulated genes were designated as hub genes.

### mRNA Expression and Survival Analysis of Hub Genes

This study explored the survival rates and associations between key genes in colorectal cancer (CRC) patients by leveraging several databases, including UALCAN (http://ualcan.path.uab.edu/), GEPIA (http://gepia.cancerpku.cn/)(28), and KM plotter (https://kmplot.com/analysis/). Survival analysis was performed using the Kaplan-Meier method combined with log-rank tests, with statistical significance set at p < 0.05 to highlight meaningful associations between gene expression levels and patient survival outcomes. To validate expression levels, CRC patient data from The Cancer Genome Atlas (TCGA) were analyzed. Based on transcripts per million (TPM) values, data were categorized into two groups for visualization in the GEPIA database: patients in the low/medium expression group had TPM values below the upper quartile, while those in the high expression group had TPM values above the upper quartile.

### Prognostic Characteristics of Hub Genes

To assess the prognostic role of hub genes, mRNA expression data from early and advanced stages of colorectal cancer (CRC) were analyzed using the GEPIA database (http://gepia.cancer-pku.cn/). Additionally, the metastatic potential of these hub genes was evaluated through the UALCAN database (http://ualcan.path.uab.edu/)(29). The cBio Cancer Genomics Portal (http://cbioportal.org)provided mutation and expression profiles of hub genes from colorectal tissue samples in the TCGA dataset. This portal offers in-depth mutation analysis, identifying gene mutations, amplifications, and deletions across 20 CRC studies. Protein levels of hub genes in tumor tissues were further examined using the Human Protein Atlas (HPA) database (https://www.proteinatlas.org/)(30), which provides immunohistochemistry-based expression data, facilitating the analysis of protein expression patterns across various human tissues.

## Results

### Identification of DEGs

Here, we studied the gene expression of the GSE164191 dataset, which contains information on 59 colorectal cancer tissues and 62 normal tissues. Significant differential gene expression was determined based on a p-value < 0.05 and |logFC| > 1. Overall, we identified 191 dysregulated genes, of which 89 were upregulated and 56 were downregulated. As shown in the volcano plot (Figure 1), the red dots represent upregulated genes, while the green dots represent downregulated genes. The complete list of upregulated and downregulated genes is provided in Table 1.

**Figure 1.**
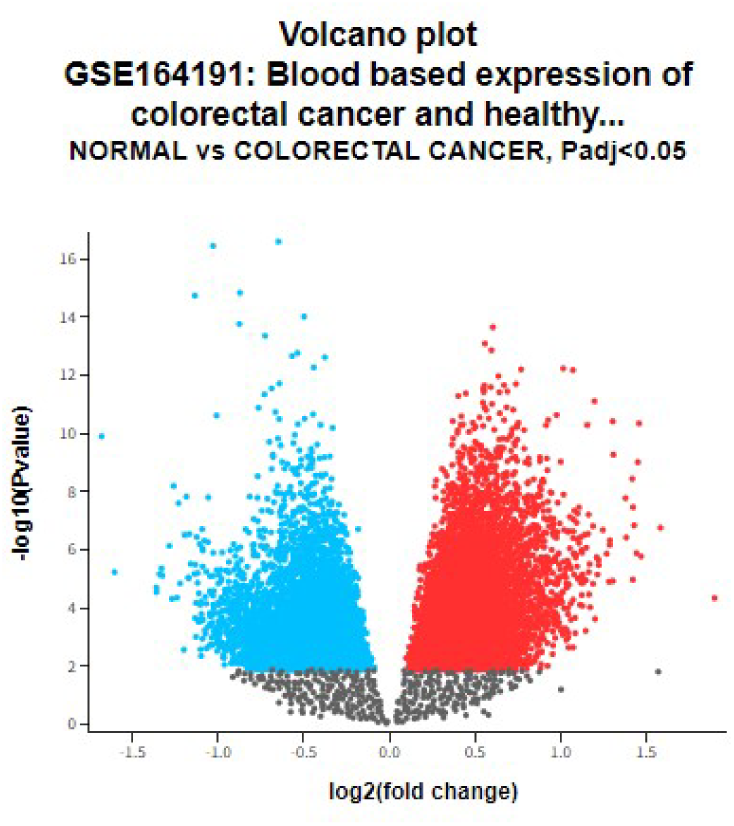
Volcano Plot of Differential Gene Expression in Colorectal Cancer vs. Normal Tissues (GSE164191 Dataset)

**Table 1:** List of Upregulated and Downregulated Differentially Expressed Genes (DEGs) in CRC.

| DEGs | Gene |
| --- | --- |
| Upregulated<br>(89) | GOLGA8A, MYBL1, TNFRSF25, SUPT16H, RBM8A, STK26, PTPN4, ZNF677, ELK4, ZBTB10, ENPP4, PIGL, GRAMD1C, SENP7, ZBTB1, CXCL8, CBLB, TMEM267, MBLAC2, ZNF540, KLRC4, TSEN15, SCML1, ENPP5, KCTD7, RBAK, TTN, MAT2B, ZNF91, ENPP4, MDFIC, TMEM263, LMAN1, ELP2, ERCC6L2, AGBL3, RABEP1, MIR146A, CCDC65, ICOS, TAF9B, LMAN1, TBC1D32, CENPK, ZNF397, E2F5, FAM188A, RDX, B4GALT6, WDR60, SH2D1A, ZEB1, ZC2HC1A, MKLN1, CCDC82, MIER3, ABCD2, ZNF770, TRIM23, S1PR5, PGBD1, HLA-DRB4, VPS13B, ARMCX4, TMEM117, ZNF883, GEN1, ARAF, UTP23, PFDN4, BRMS1L, B3GALT2, CD22, ZNF404, CXorf57, ZNF91, THAP9, FSD1L, C9orf3, N4BP2L1, RGS1, HLA-DRB4. |
| Downregulated<br>(56) | PPP3R1, C1QC, TP53I11, SERPINA3, MMP9, GRPR, C1QB, TBL1X, IRS1, GPER1, PNPLA1, MYO7B, USF1, MAGI1, NFIC, ERFE, FOXC1, LYPD8, UNC5D, PGLYRP1, OLAH, MYL9, PCYT1B, LMO2, ZNF704, NLRP3, CD177, SCGB2A2, RNF130, GRB10, CPNE4, PNLIPRP2, PRKAA2, ERC2, NDST4, SIGLEC11, EFCAB6, NR2F2, DCT, ARHGEF12, ACSM2A, ZNF536, MCF2, DSC3, ONECUT2, TMEM150C, PCDHB6, CACNA1E, ALB, NOVA1, NLGN1, CAGE1, CDH7, RAP1GAP, SLC17A1, SIM1. |

### Functional enrichment analysis of DEGs

DAVID software was used to analyze a total of 89 upregulated and 56 downregulated genes, revealing the top five significant terms from the GO enrichment analysis across different categories. In the Biological Process (BP) category, upregulated DEGs were involved in transcription regulation by RNA polymerase II, immune response, negative regulation of transcription by RNA polymerase II, DNA-templated transcription, and cell adhesion molecule production (Fig. 2a), while downregulated genes were associated with calcium-dependent cell-cell adhesion via plasma membrane adhesion molecules, glucose homeostasis, transcription regulation by RNA polymerase II, fatty acid biosynthesis, and innate immune response. In terms of the Cellular Component (CC) category, upregulated DEGs were primarily localized to the nucleus, nucleoplasm, and Golgi membrane (Fig. 2b), while downregulated DEGs were linked to the postsynapse, complement component C1q complex, complement component C1 complex, membrane, and blood microparticle. For the Molecular Function (MF) category, upregulated DEGs showed enrichment in RNA polymerase II-specific DNA-binding transcription factor activity, protein homodimerization, metal ion binding, and DNA-binding transcription repressor activity (Fig. 2c), whereas downregulated DEGs were related to calcium ion binding, identical protein binding, and sequence-specific DNA binding. Additionally, KEGG pathway enrichment analysis indicated that downregulated DEGs were involved in axon guidance, the Pertussis pathway, and the C-type lectin receptor signaling pathway, whereas upregulated DEGs were mainly associated with the Herpes simplex virus 1 infection pathway.

**Figure 2.**
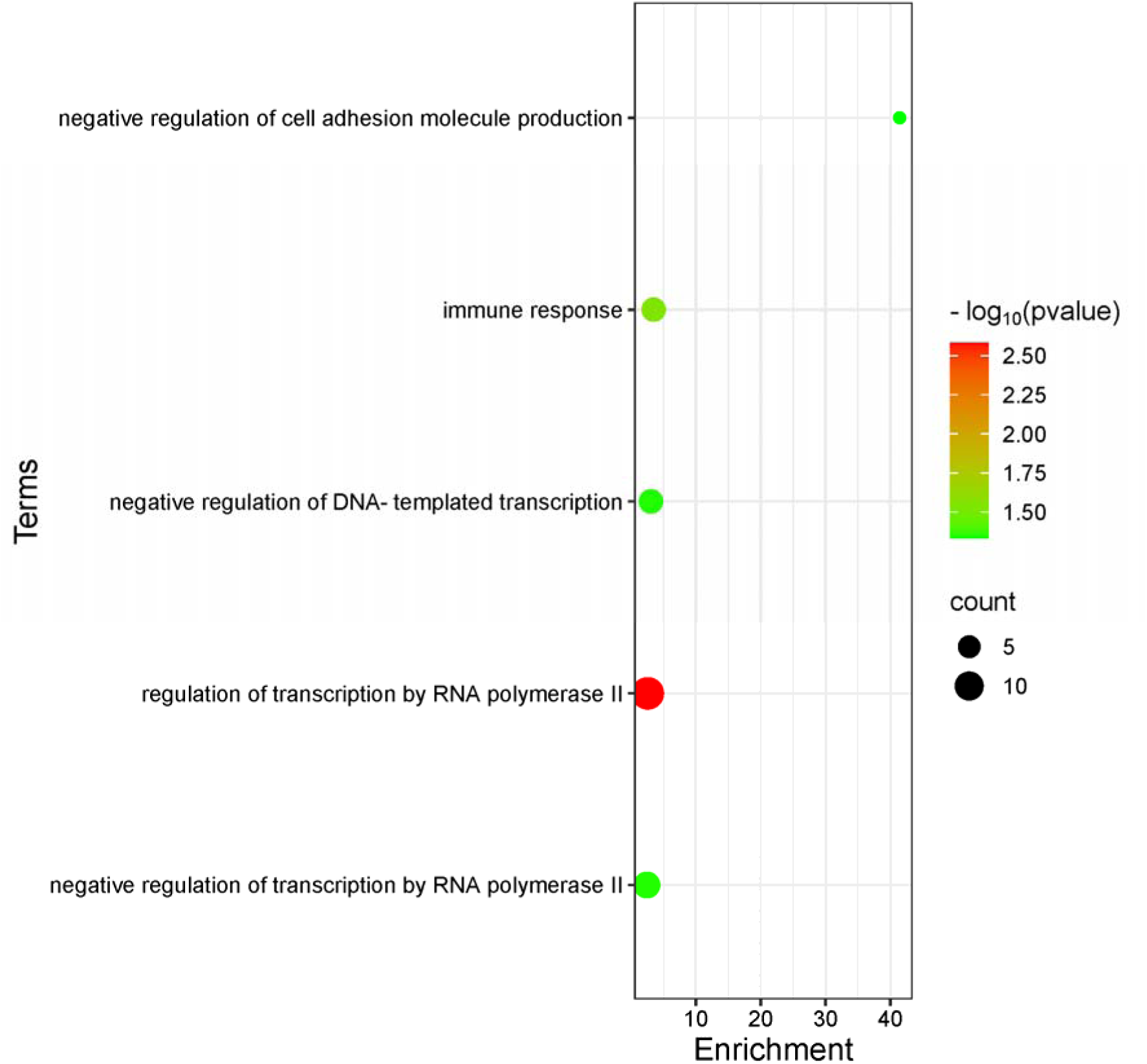
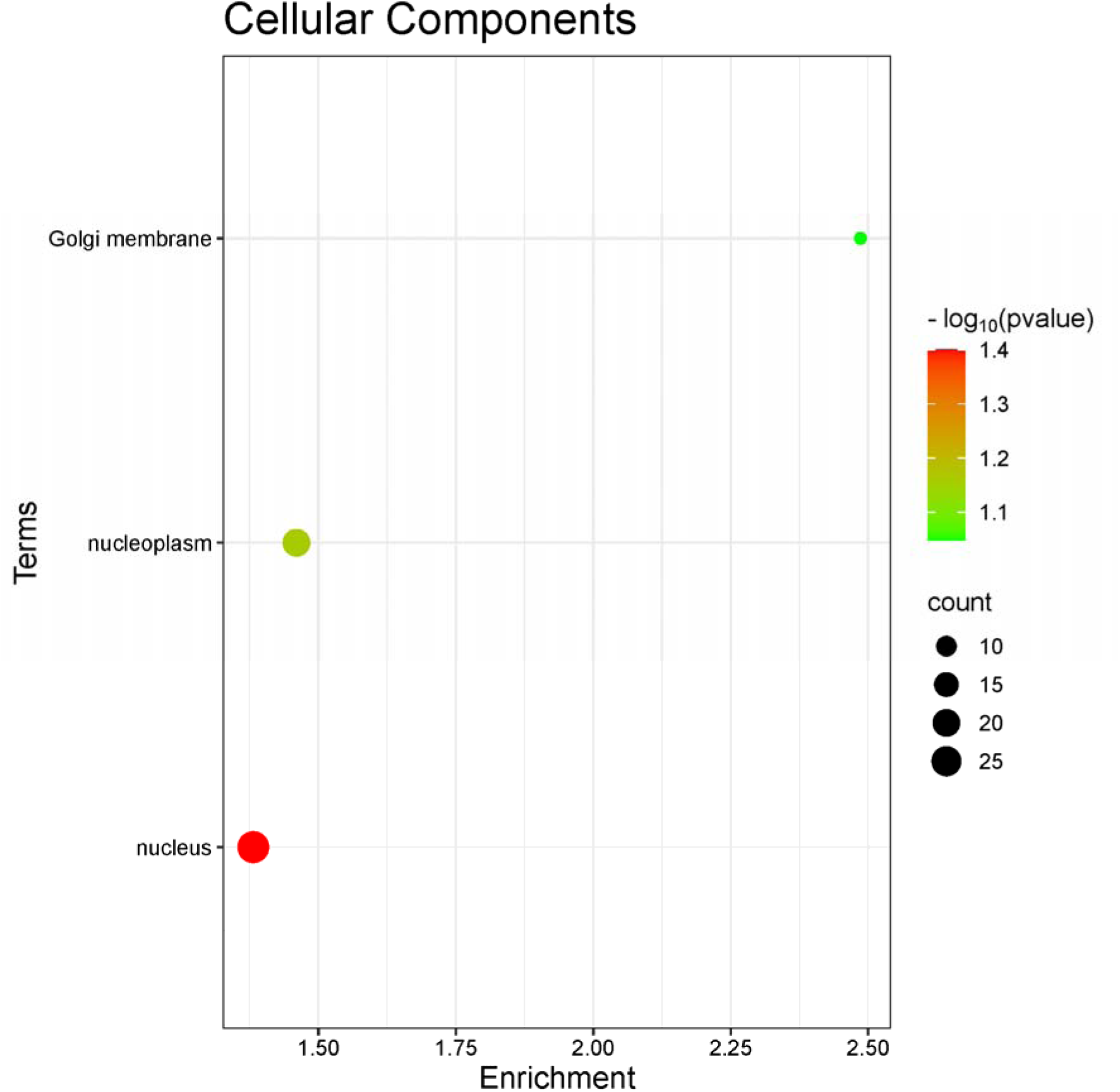
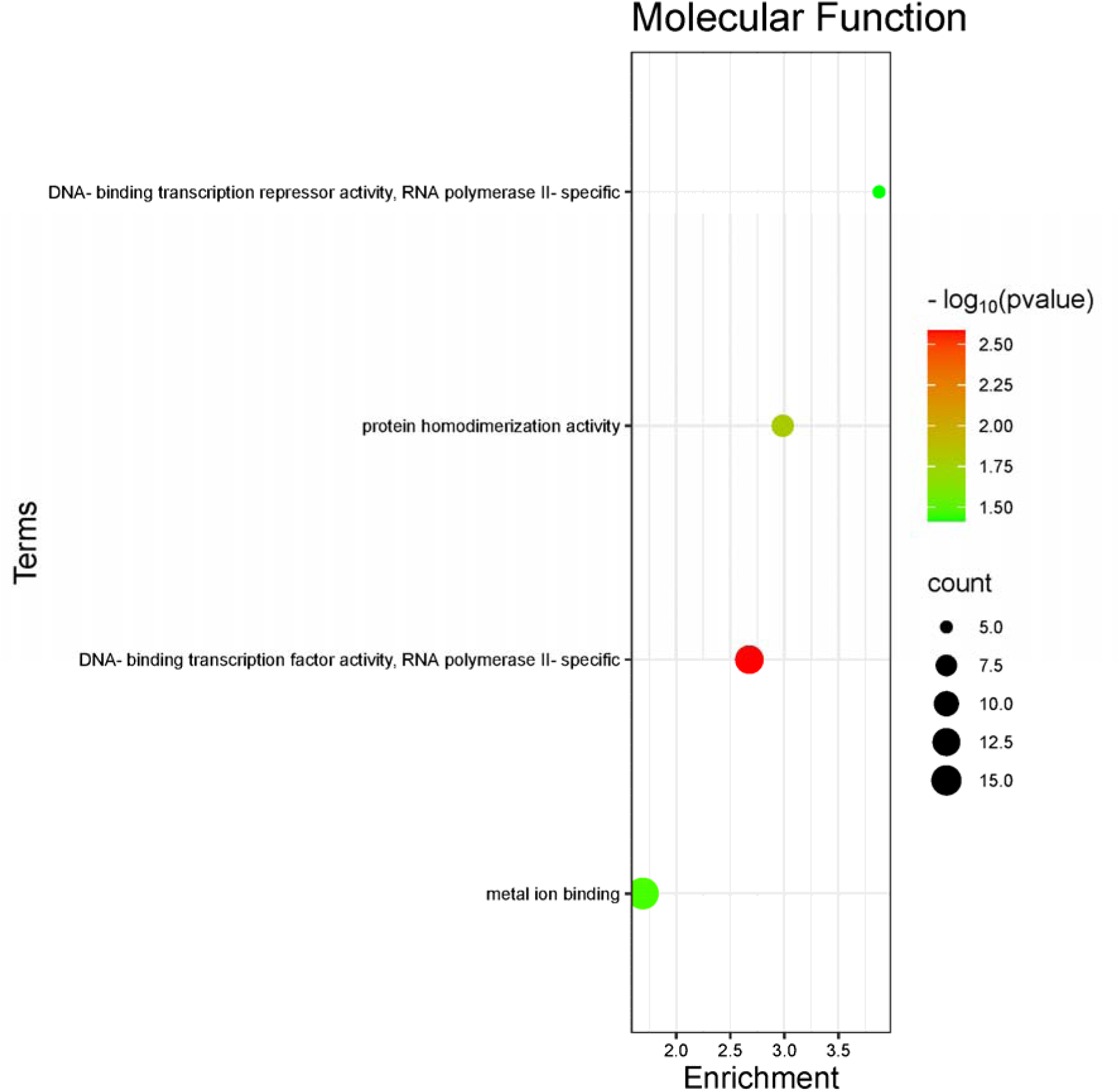
Bubble map illustrating GO and KEGG pathway analyses for upregulated DEGs. The top 5 terms from GO and KEGG pathway enrichment analyses. Statistical significance was set at a p-value < 0.05. Categories include: (a) Biological Processes, (b) Cellular Components, (c) Molecular Function.

### PI network construction and module analysis

The PPI network of DEGs in colorectal cancer was constructed using data from the STRING database. When 145 DEGs were uploaded, a total of 168 genes were mapped into the PPI network, which included 131 nodes and 52 edges, with a PPI enrichment p-value of 0.21 (Fig. 3a). Additionally, an interaction network of the 145 DEGs and their neighboring genes was built using FunRich (Fig. 3b). Significant modules within the network were identified using the MCODE plugin, with the top two functional clusters selected (module 1, MCODE score = 4.50; module 2, MCODE score = 3.00) (Fig. 3c).

**Figure 3.**
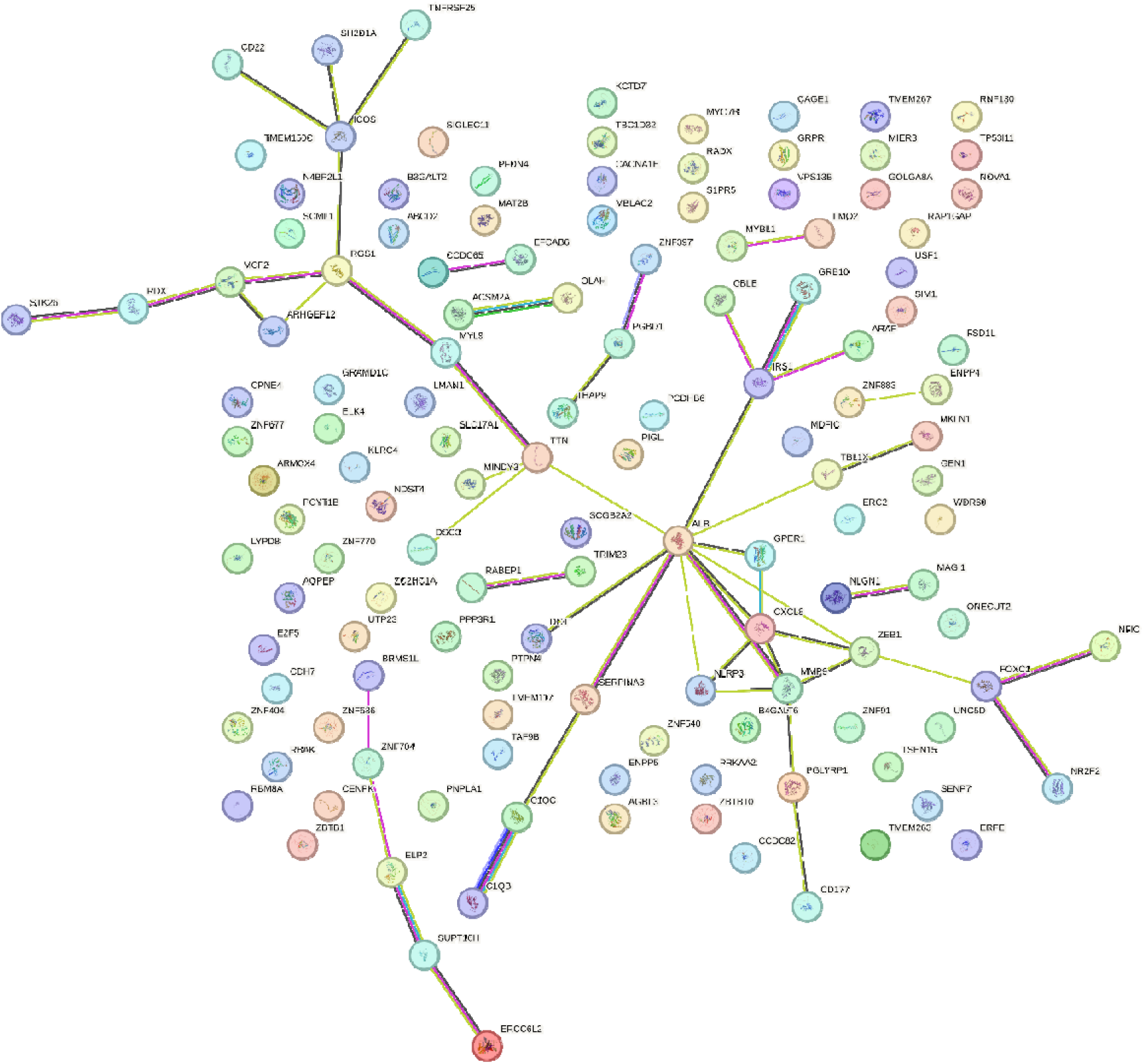
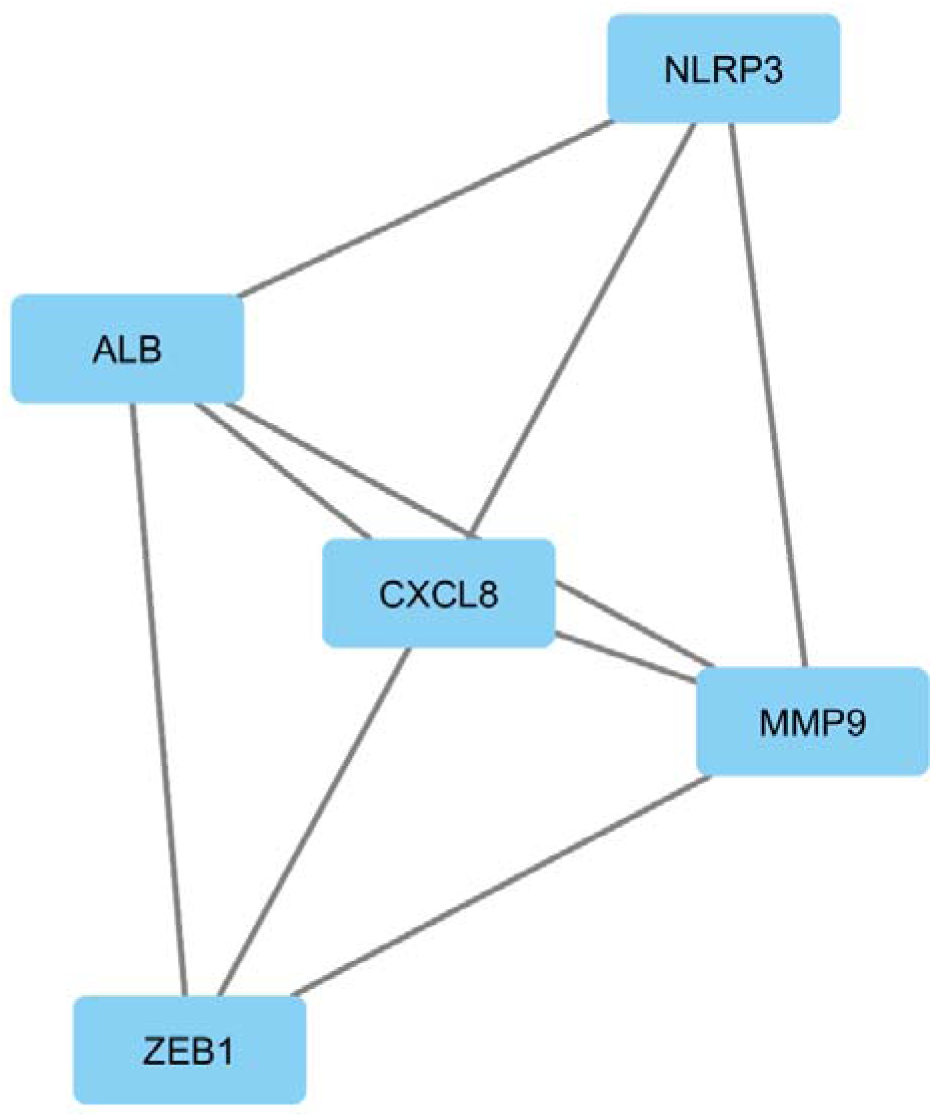
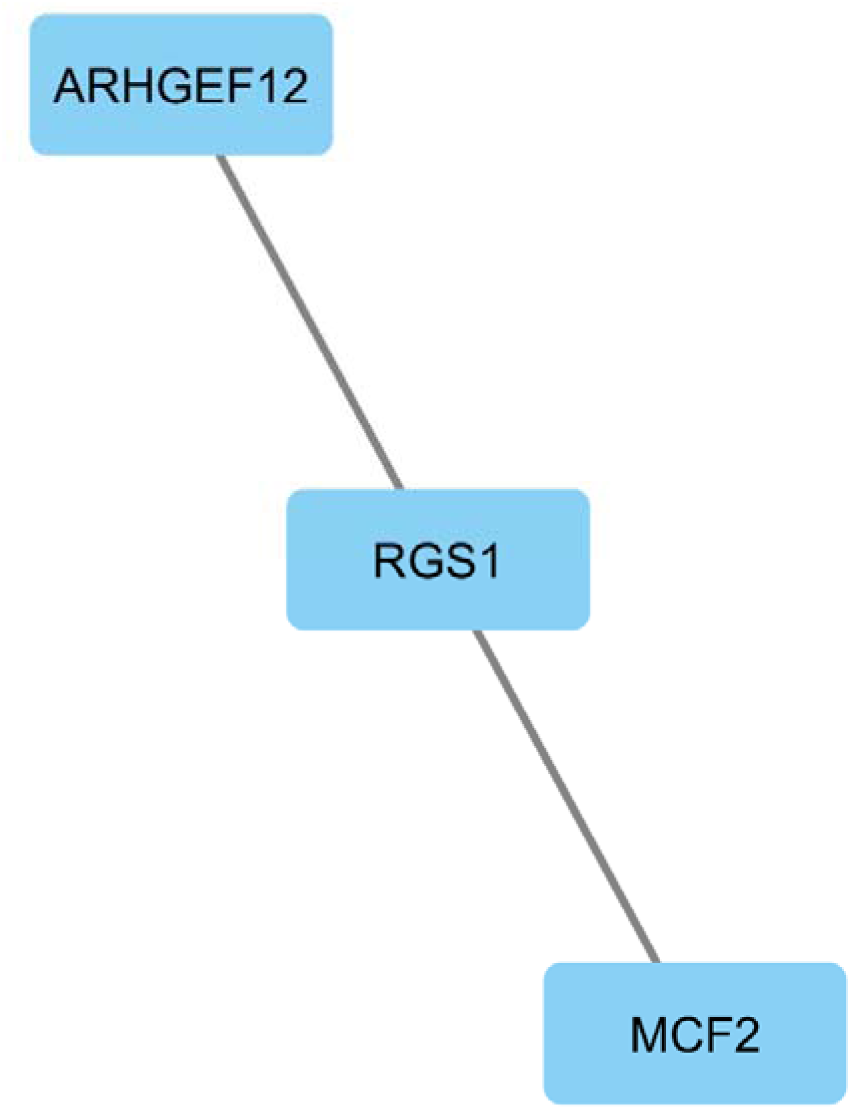
Protein-protein interaction (PPI) network analysis and module identification of differentially expressed genes (DEGs) in colorectal cancer (CRC). (a) The PPI network, constructed using the STRING database, displays DEGs with a total of 131 nodes and 52 edges Protein-protein interaction (PPI) network analysis and module identification of differentially expressed genes (DEGs) in colorectal cancer (CRC). The two most significant modules were identified using the MCODE plugin in Cytoscape. (a) Module 1 with a score of 4.50. Protein-protein interaction (PPI) network analysis and module identification of differentially expressed genes (DEGs) in colorectal cancer (CRC). The two most significant modules were identified using the MCODE plugin in Cytoscape. (b) Module 2 with a score of 3.00.

Using the cytoHubba plugin, ten genes with the highest degree scores (MMP9, IRS1, FOXC1, CXCL8, TTN, RGS1, ICOS, ALB, MCF2, and ZEB1) were identified as hub genes for colorectal cancer(Fig. 4a). All of these hub genes were found to be upregulated DEGs, as shown in Table 2. The STRING database was then employed to construct the PPI network for these hub genes (Fig. 4b), and FunRich software was used to illustrate the interaction network of the hub genes along with their related genes (Fig. 4c). In Fig. 4a, the PPI network of the hub genes contains 10 nodes and 11 edges, with an average local clustering coefficient of 0.78 and a PPI enrichment p-value of less than 1.0e−16.

**Figure 4.**
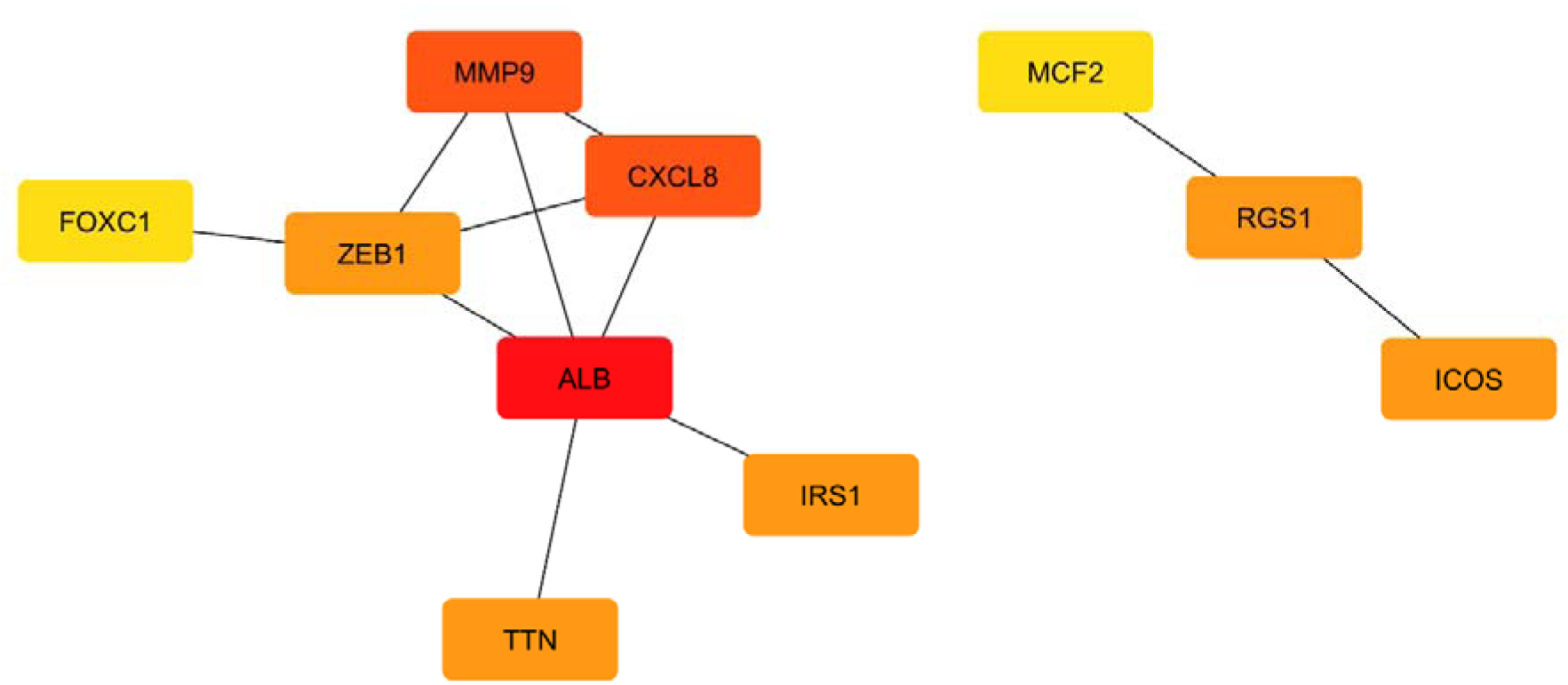
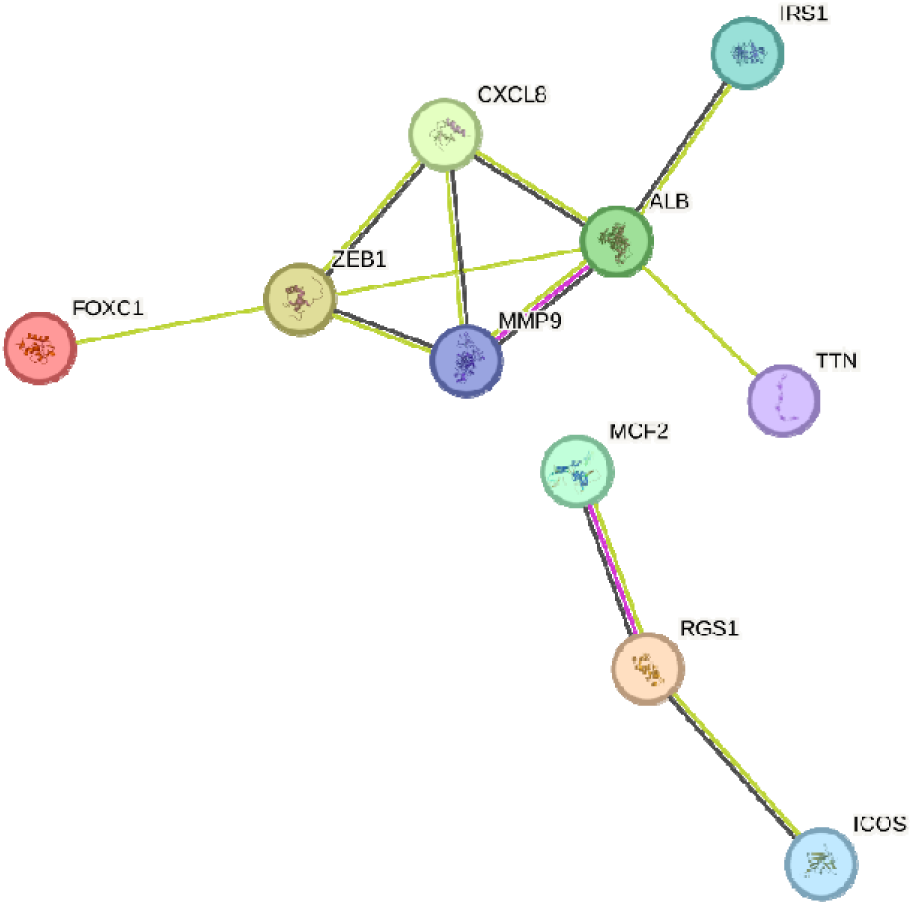
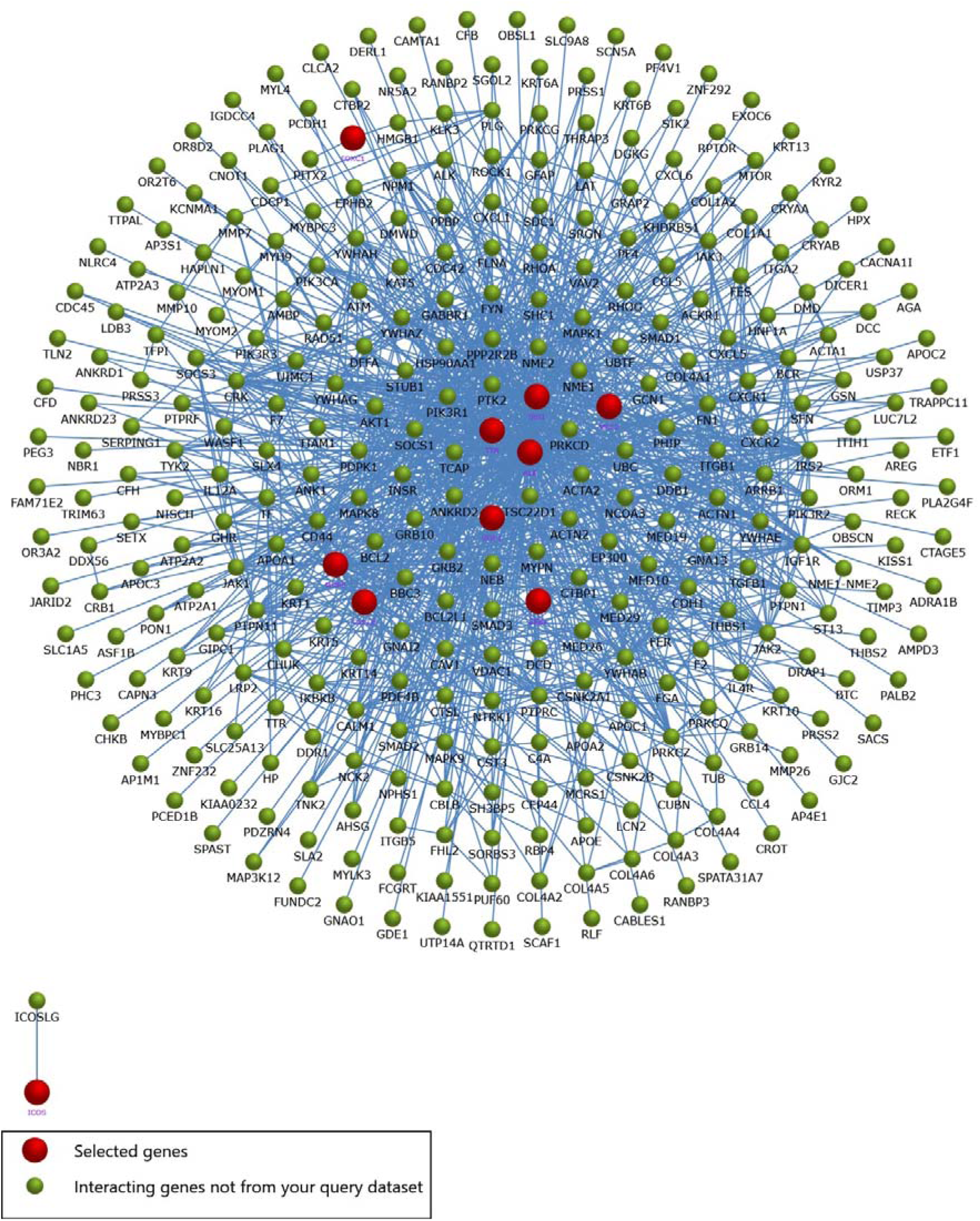
PPI network and coexpression analysis of hub genes in Colorectal Cancer (CRC). (a) The top 10 key genes identified using the Degree plug-in of CytoHubba (b) PPI network of hub genes generated using the STRING online database. (c) Interaction network of hub genes along with their neighboring genes created with FunRich software.

### Expression validation and survival analysis of hub genes in CRC

To confirm the previous findings and validate the potential hub genes identified for colorectal cancer, survival analysis was conducted using several databases, including KM plotter, GEPIA, and UALCAN. These analyses provided insights into the prognostic significance of the selected hub genes. It was identified as four genes was statistically significant ((p < .05) in colorectal cancer patients.High expression levels of CXCL8, FOXC1, ICOS, and MCF2 were associated with reduced survival rates in colorectal cancer patients (Fig.5a). Furthermore, GEPIA analysis confirmed that the expression levels of four hub genes (CXCL8, FOXC1, ICOS, and MCF2) were significantly elevated in tumor samples (Figure 5(b)).These findings further validate that the elevated expression levels of hub genes, along with the survival data from patient samples analyzed using the GEPIA database, serve as significant prognostic markers in colorectal cancer tumorigenesis.

### Analysis of prognostic values for hub genes

Understanding the role of oncogenic genes in cancer progression requires evaluating their prognostic significance in relation to cancer advancement. GEPIA database is used to validate the prognostic value of hub genes like CXCL8, FOXC1, ICOS, and MCF2, as shown in Figure 6(a). Analysis of mRNA expression in CRC datasets showed significantly elevated levels of these hub genes in advanced stages. Furthermore, the nodal metastasis status was evaluated using the UALCAN database, consistently indicating an increased metastatic potential for all hub genes, as shown in Figure 6(b). These findings collectively indicate that the hub genes possess oncogenic characteristics and are involved in the tumorigenesis of colorectal cancer (CRC).

**Figure 6.**
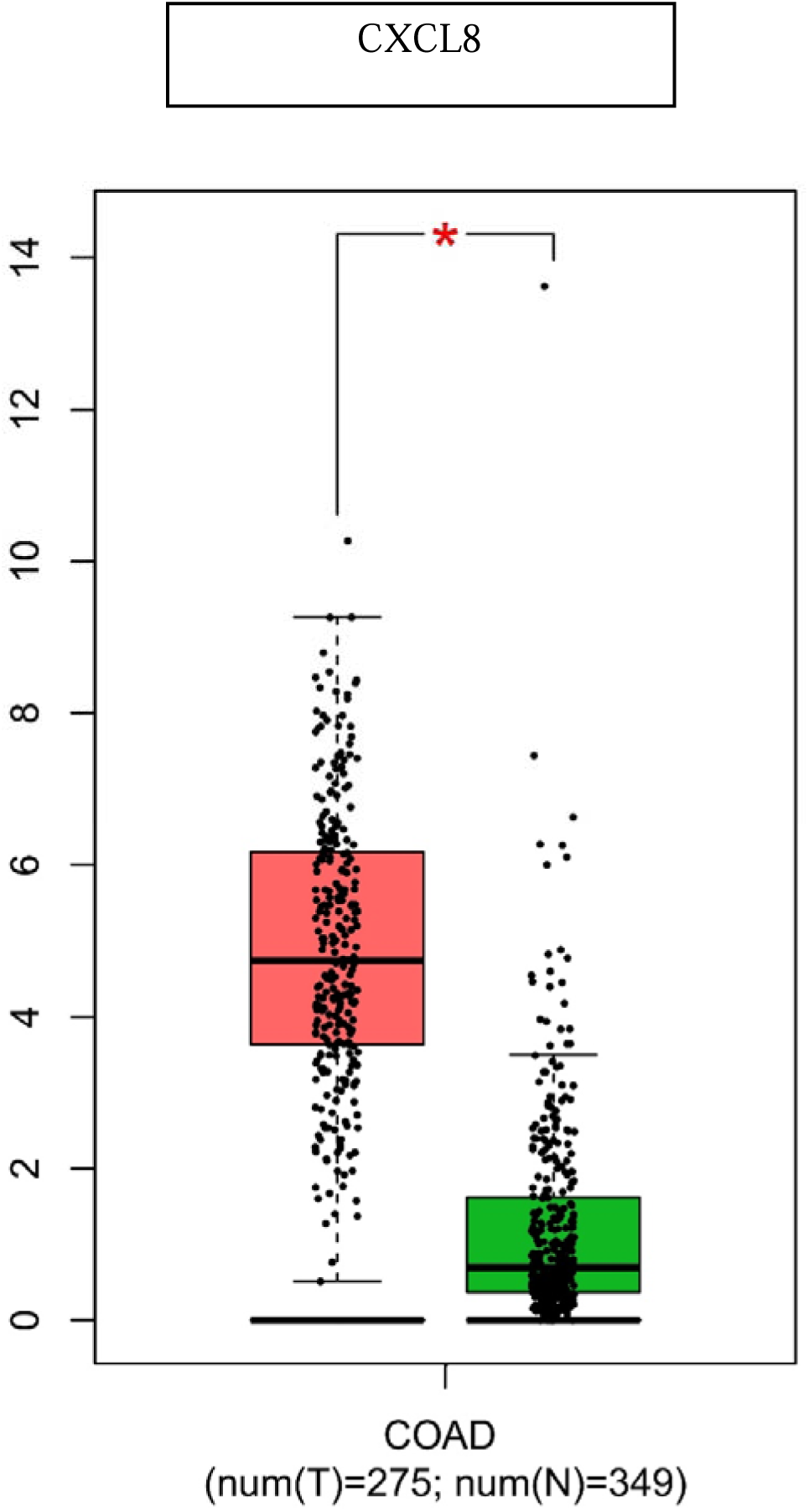
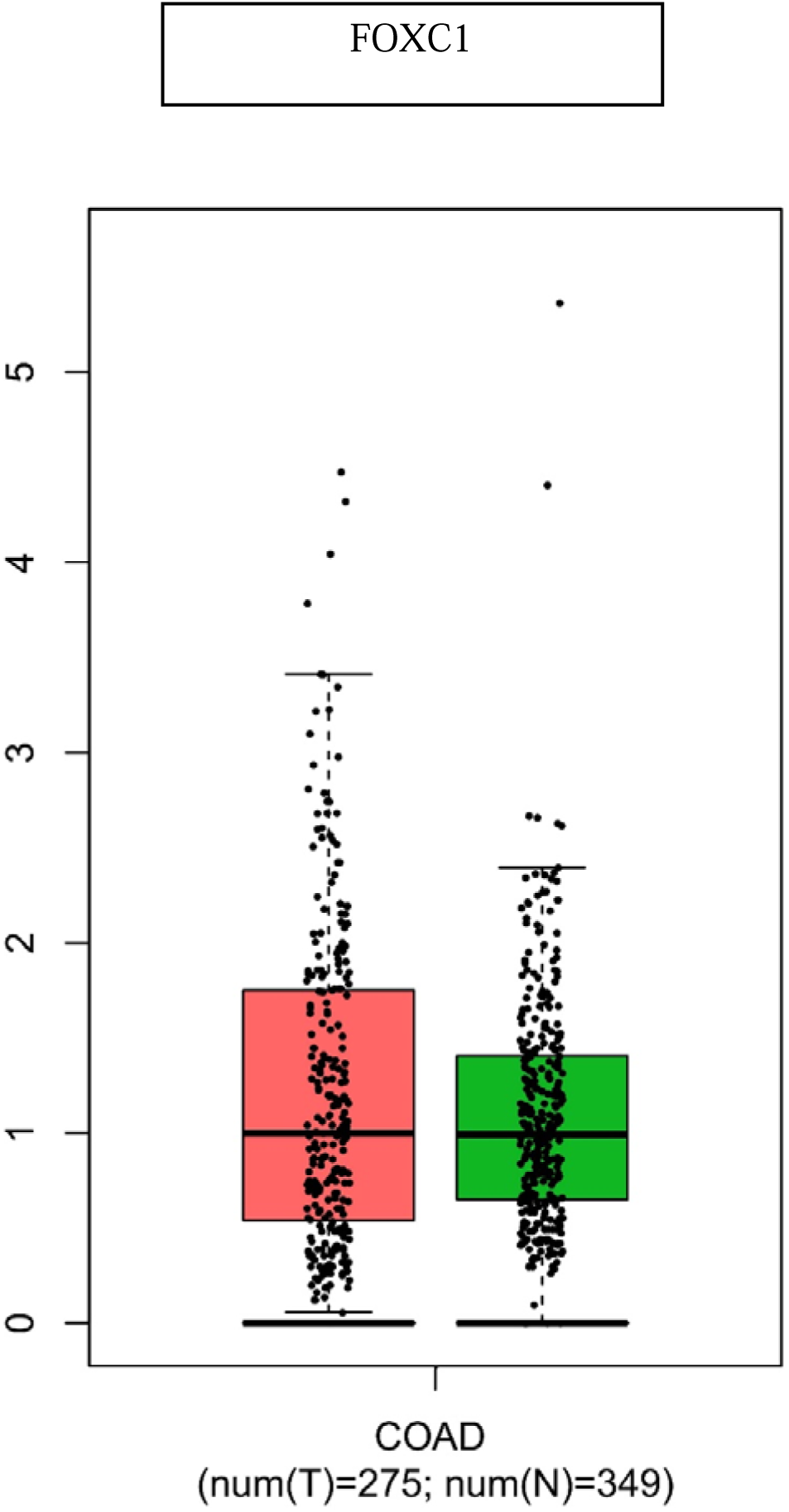
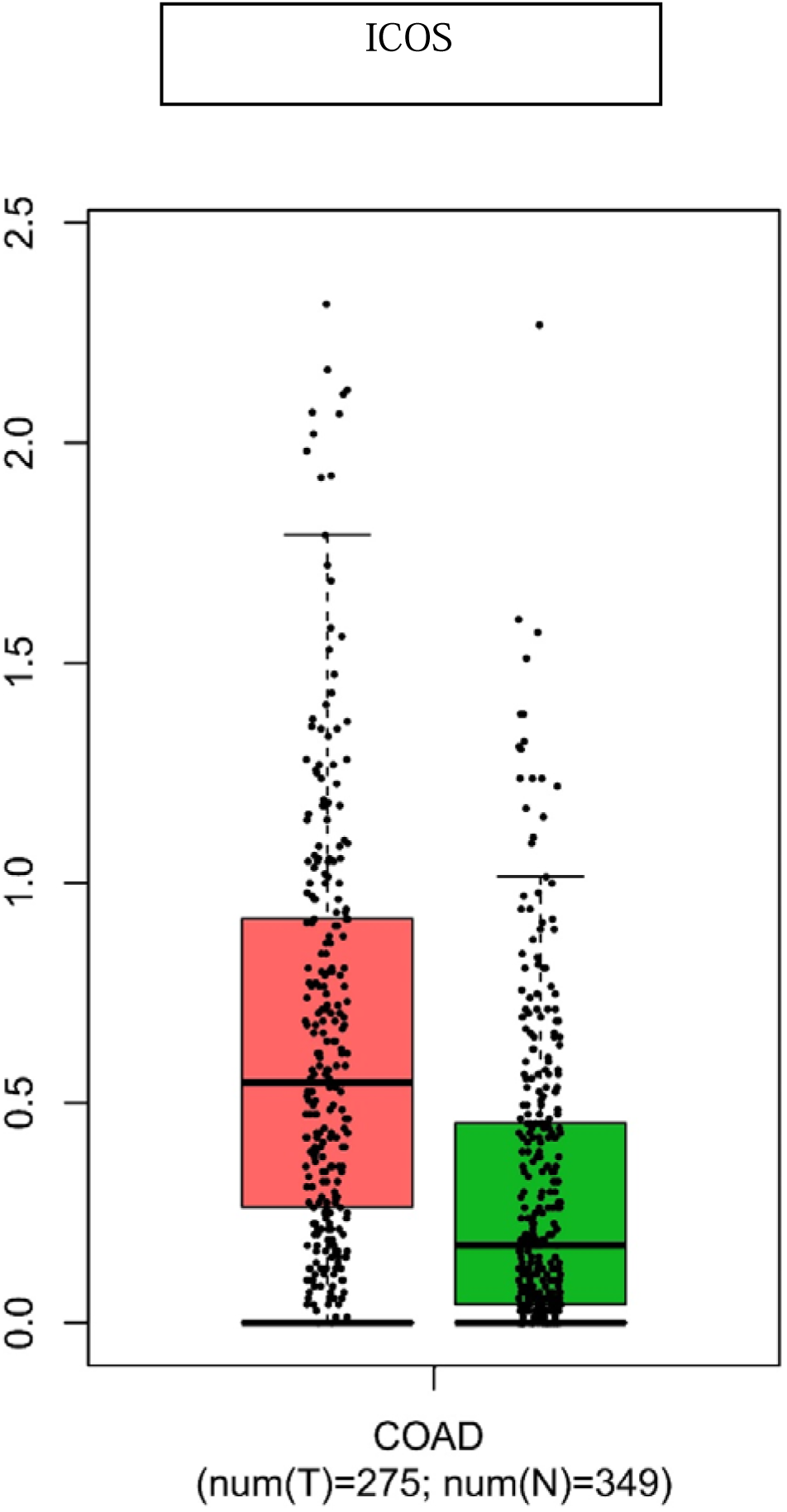
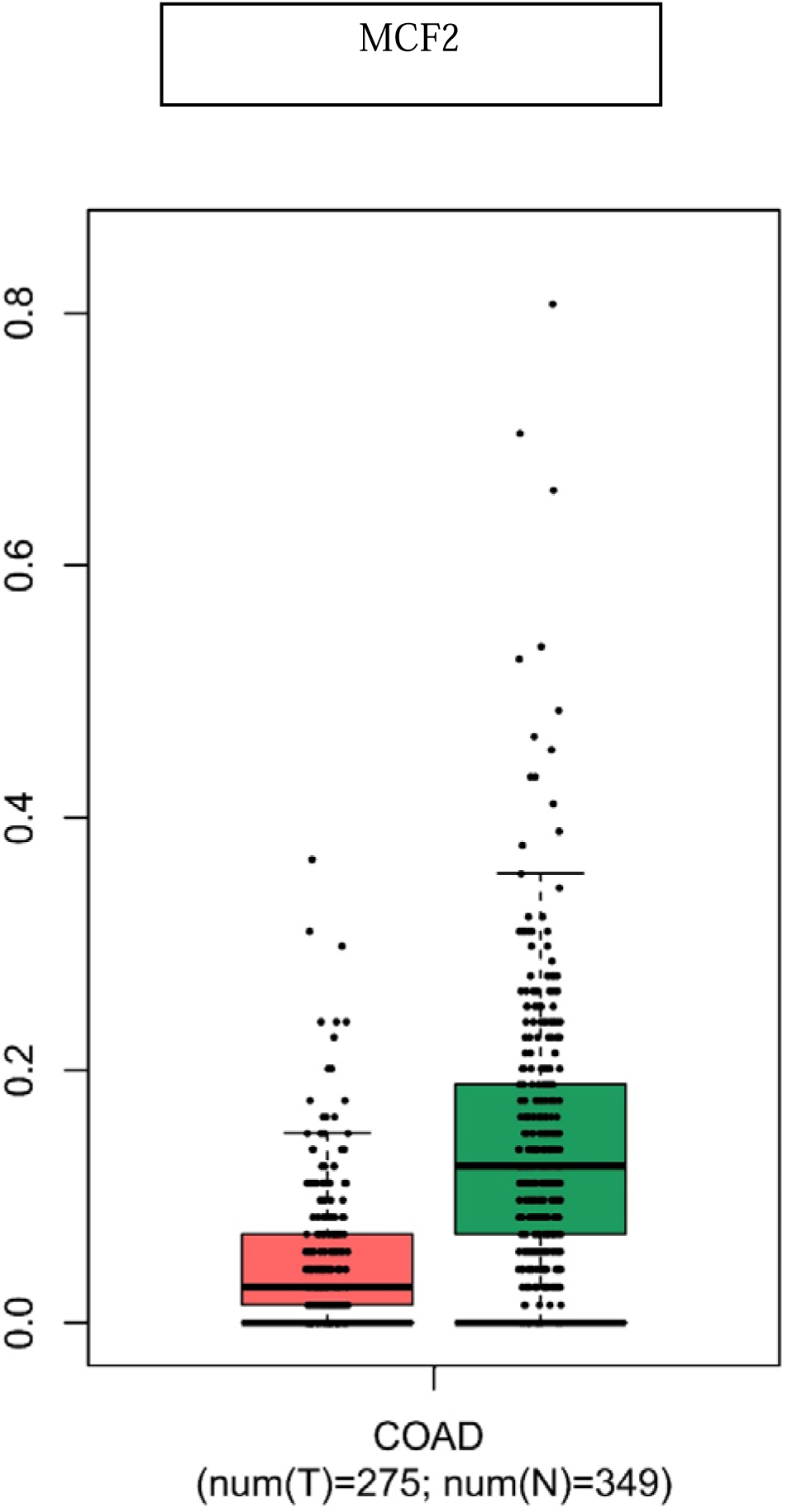
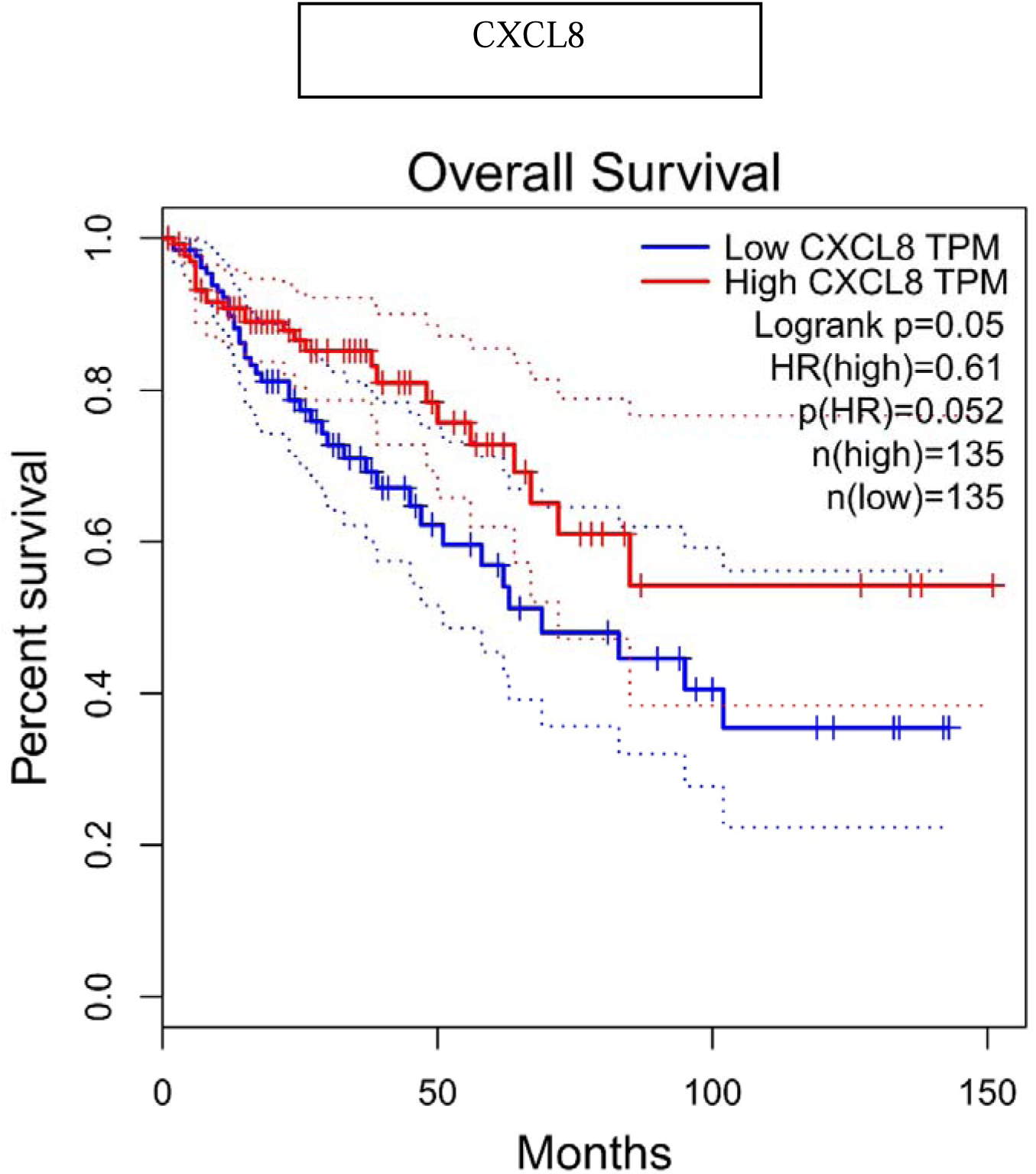
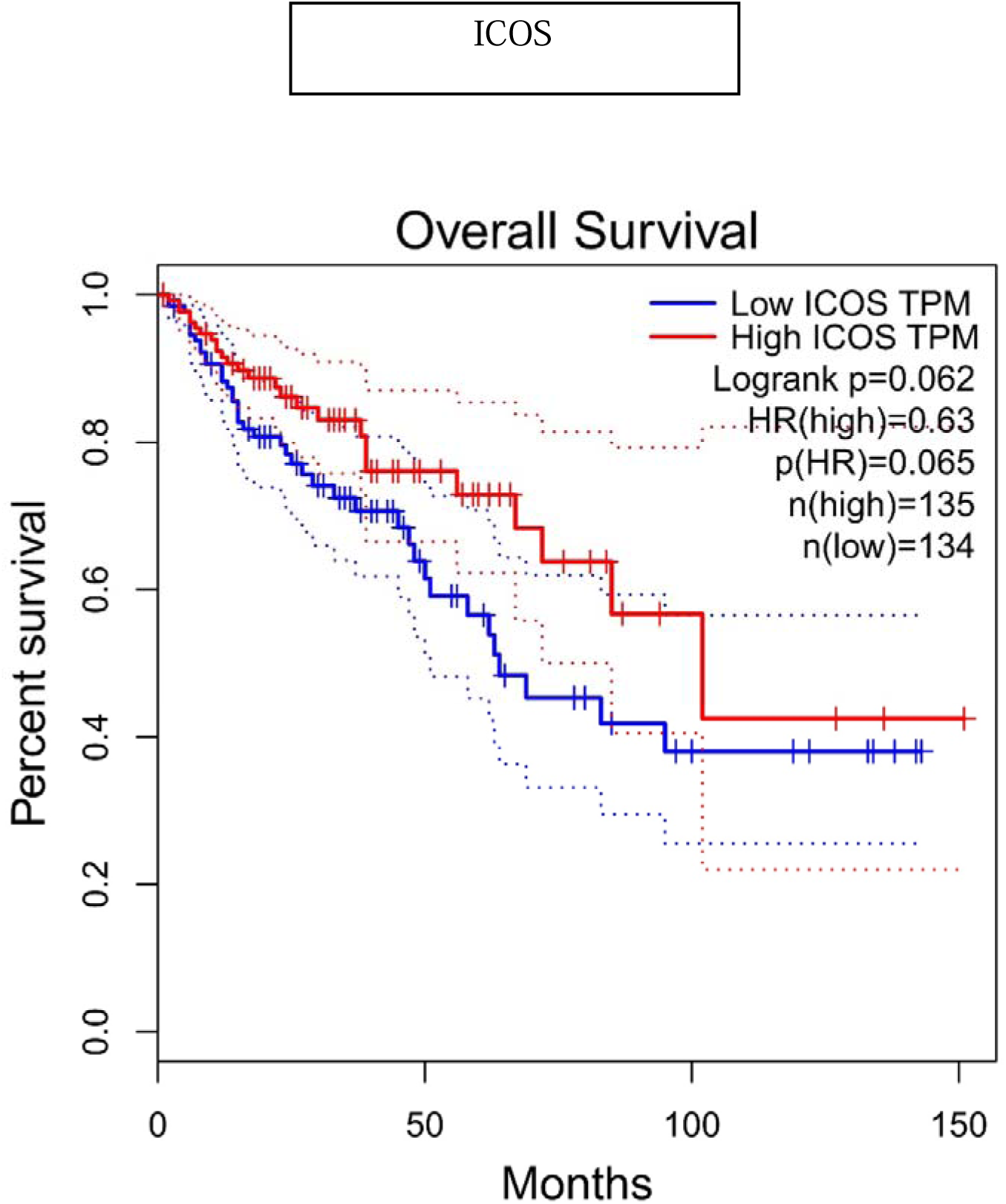
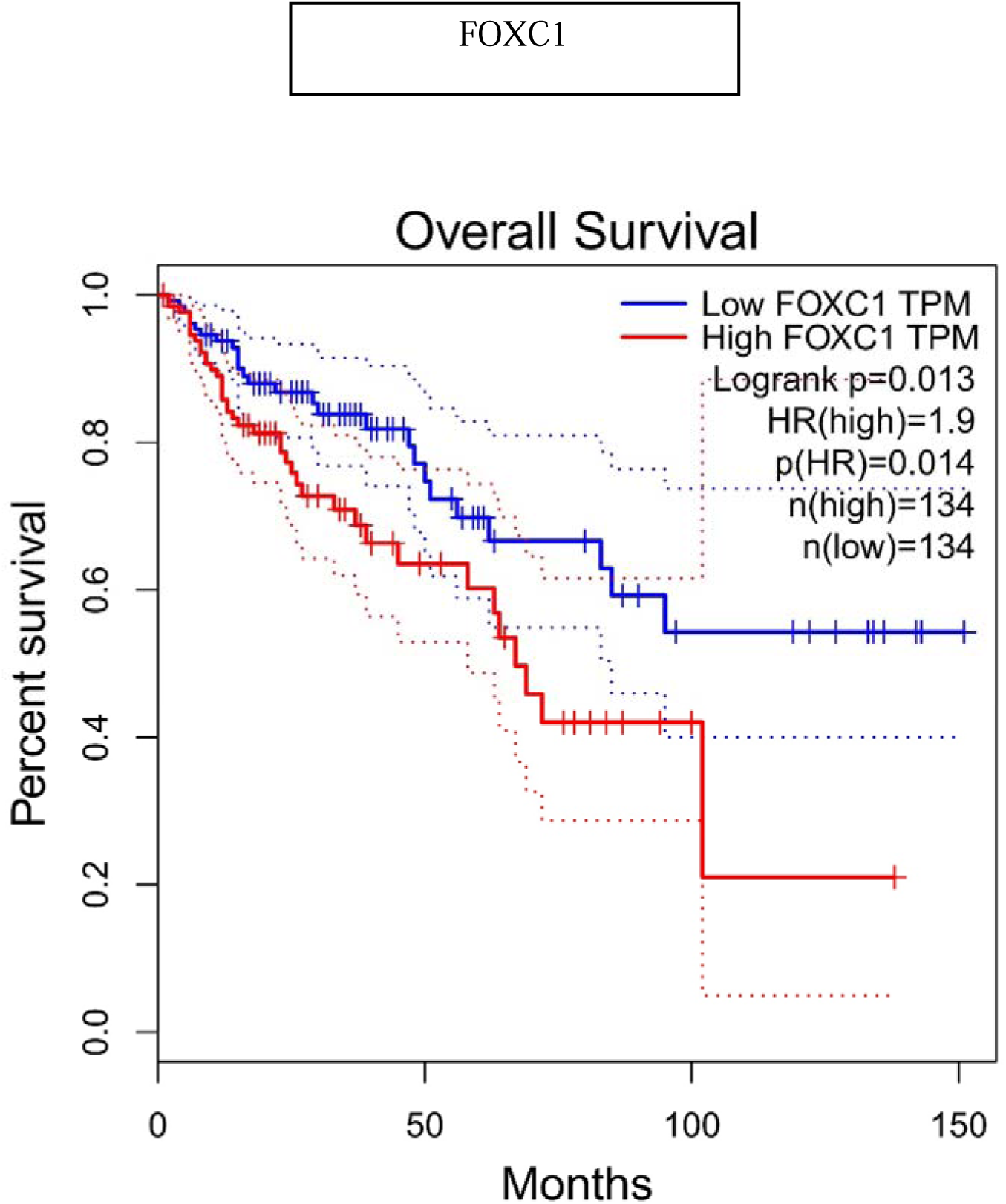
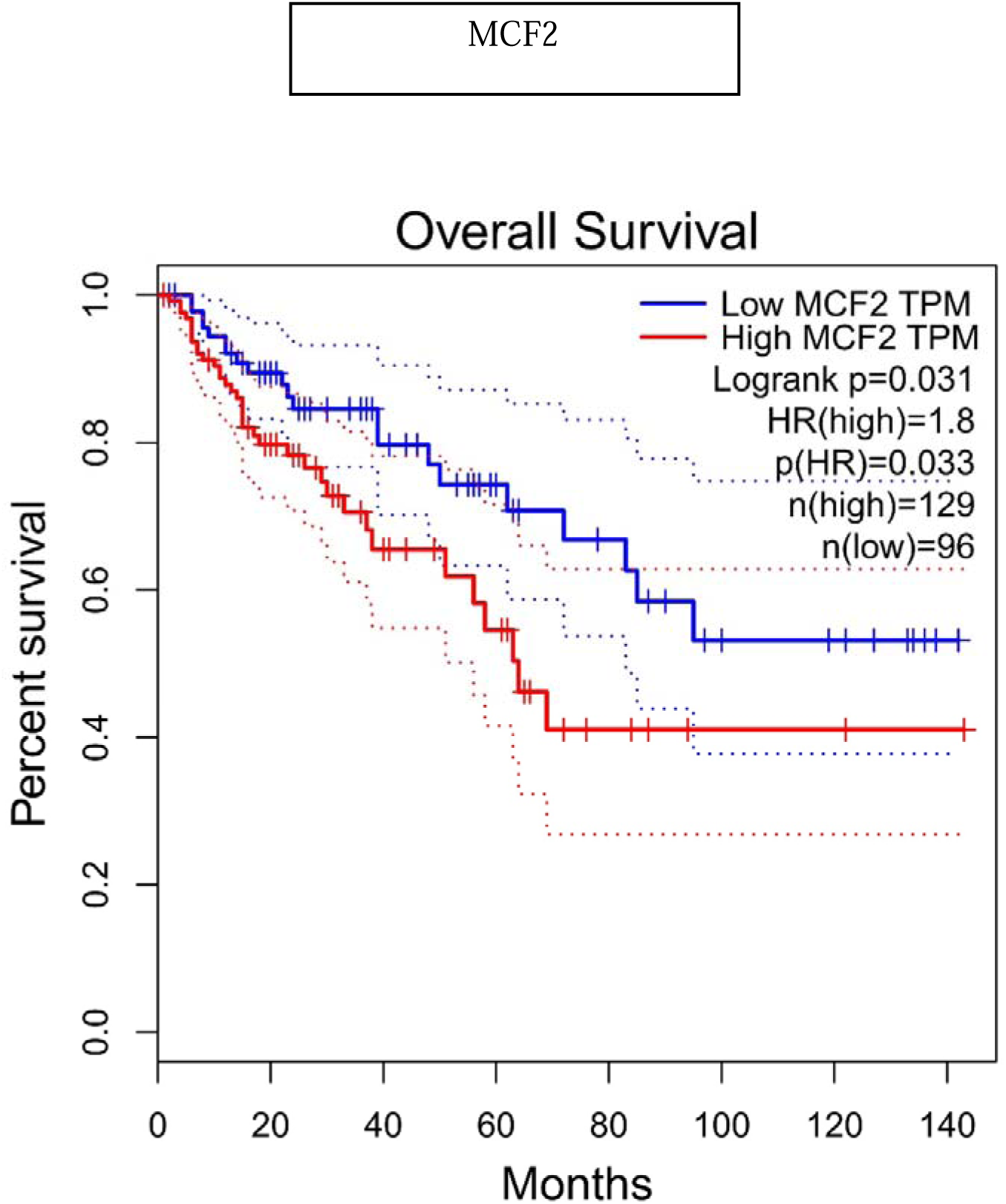
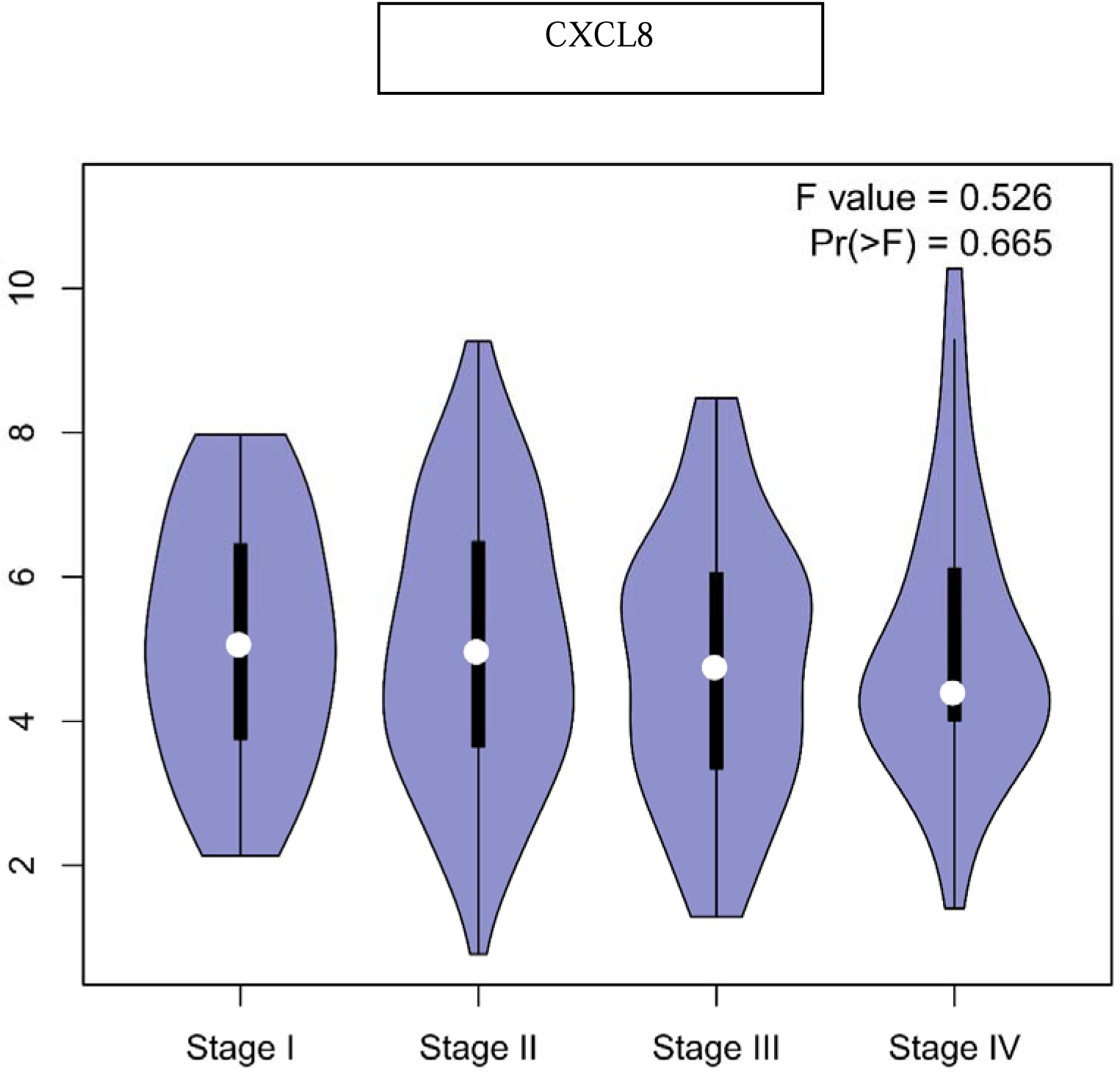
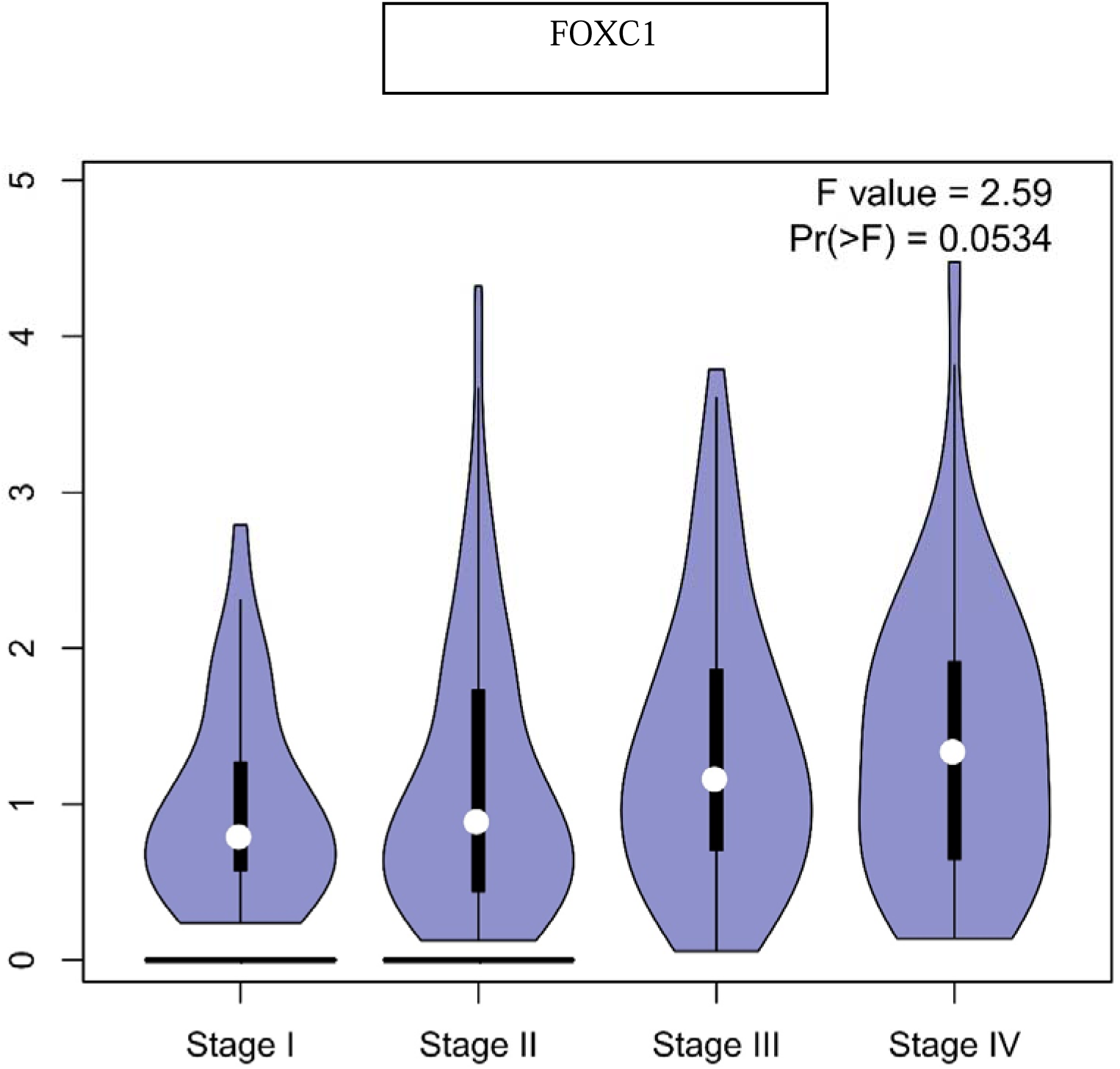
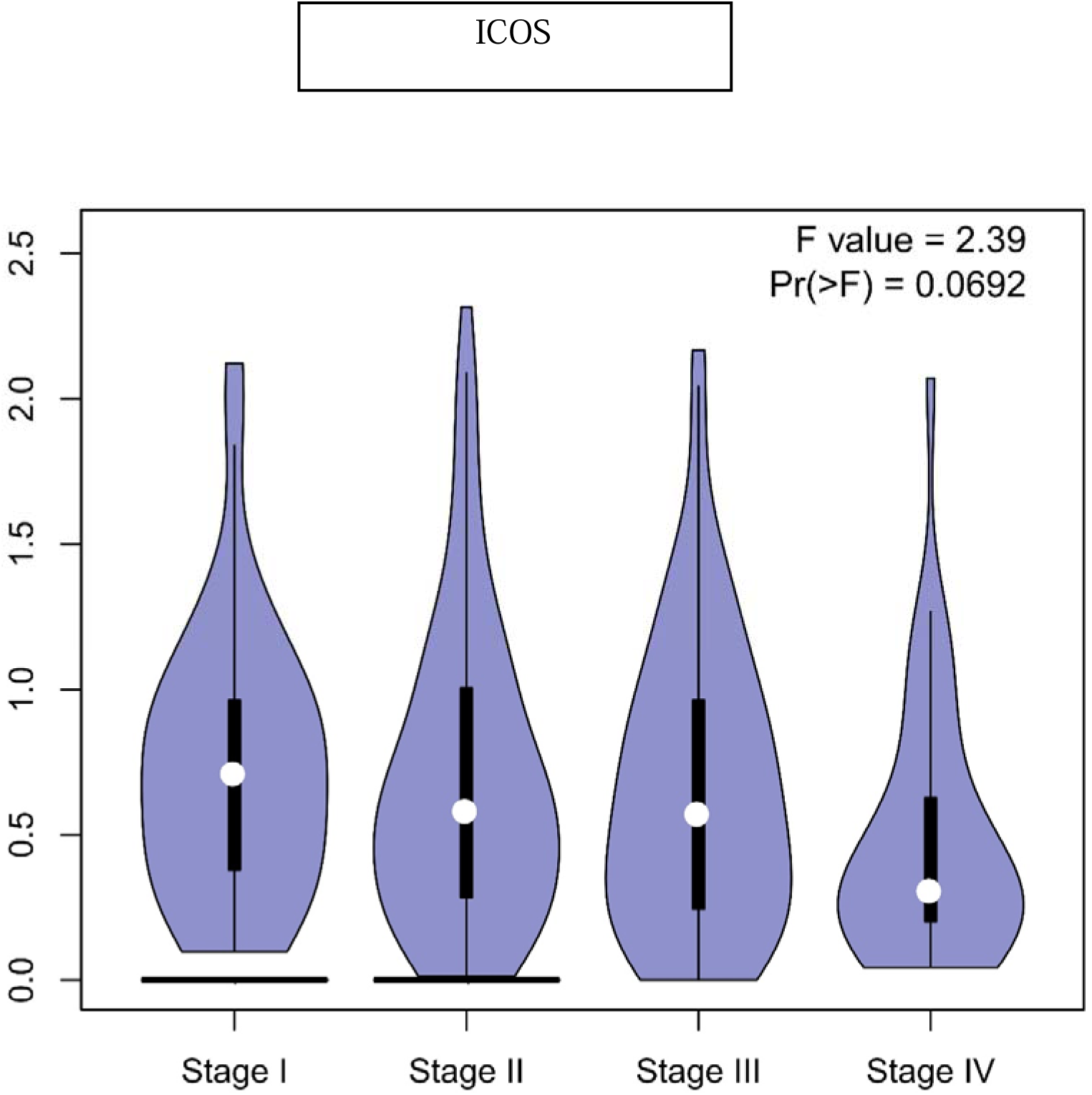
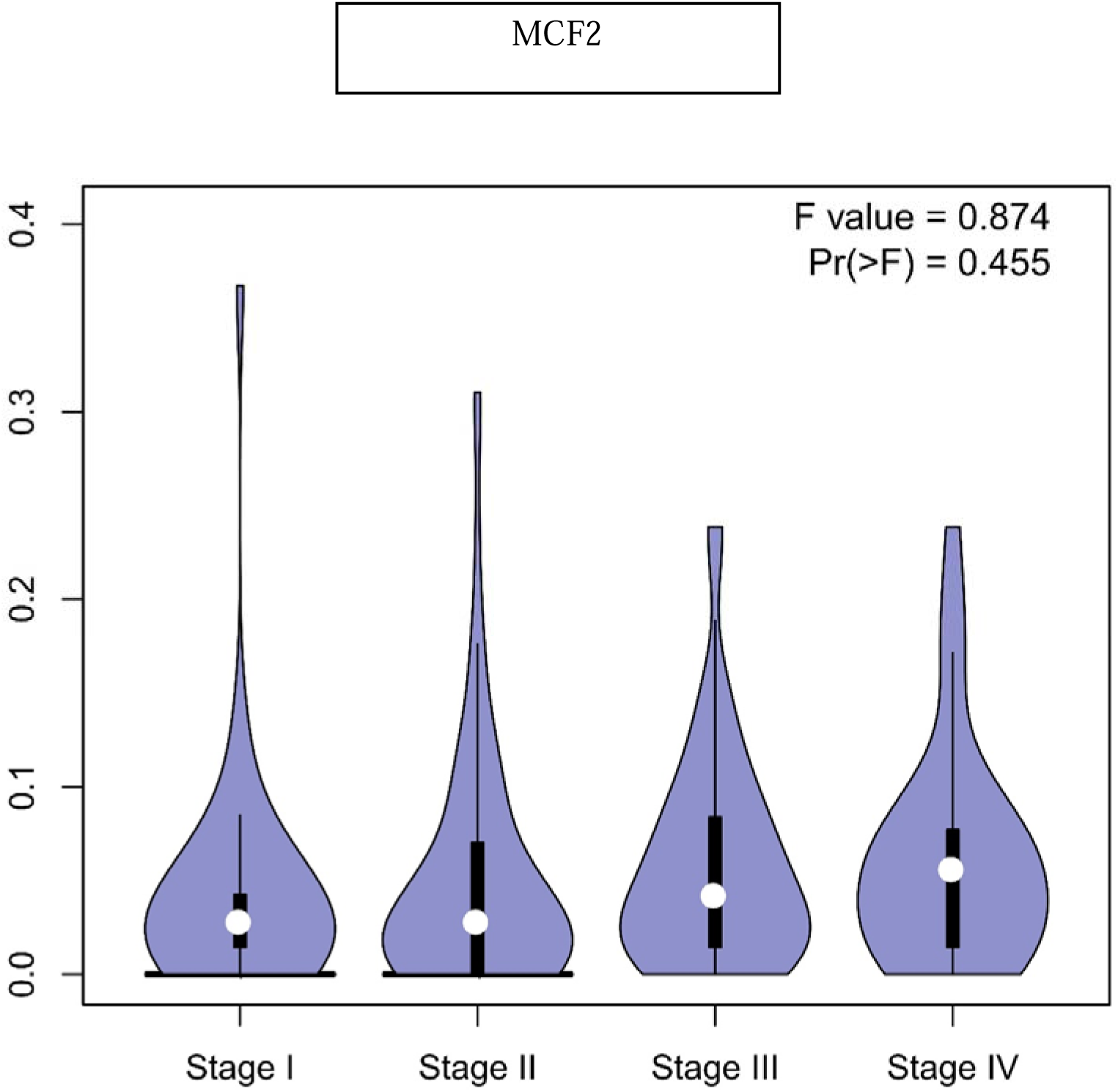
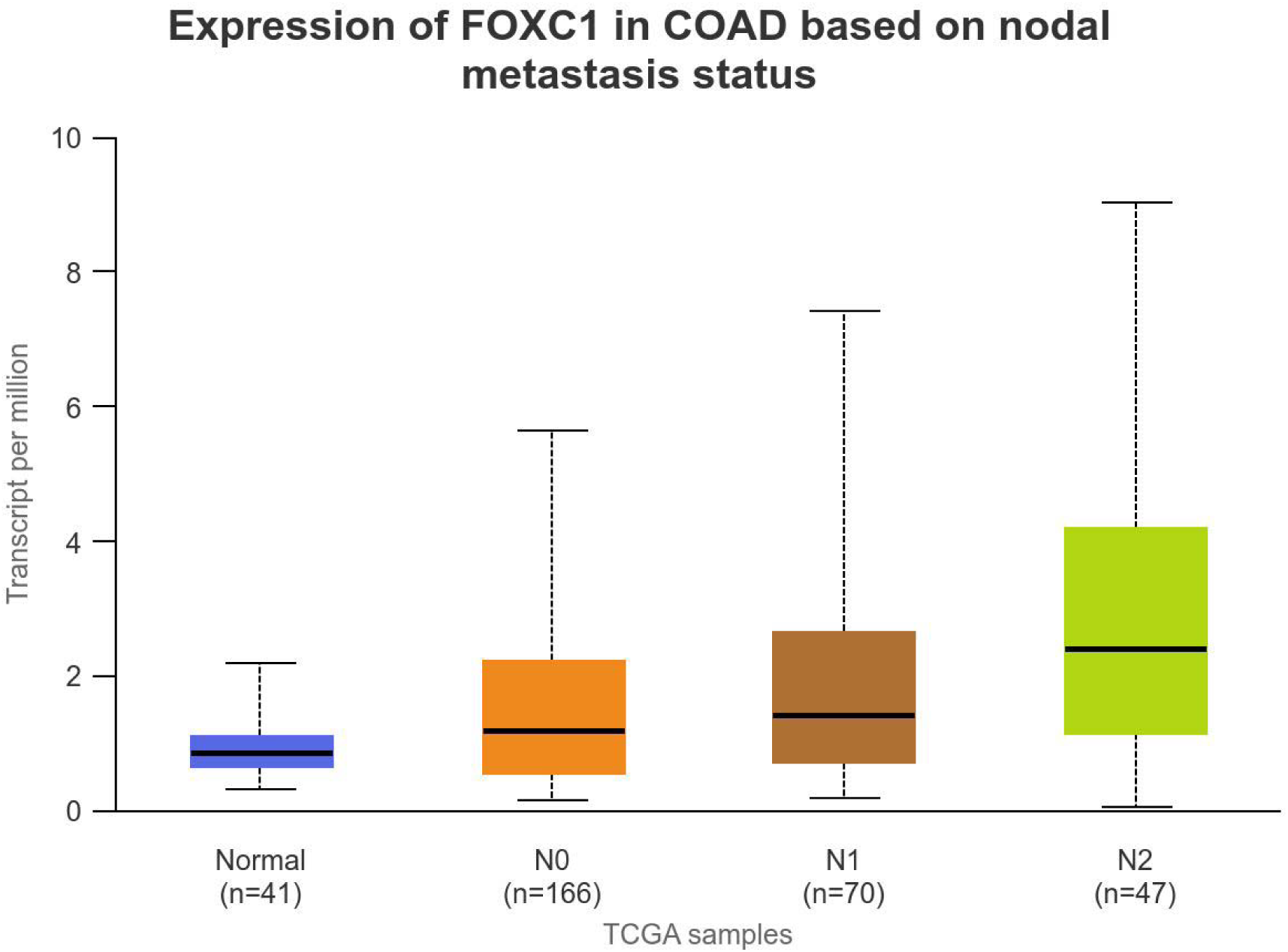
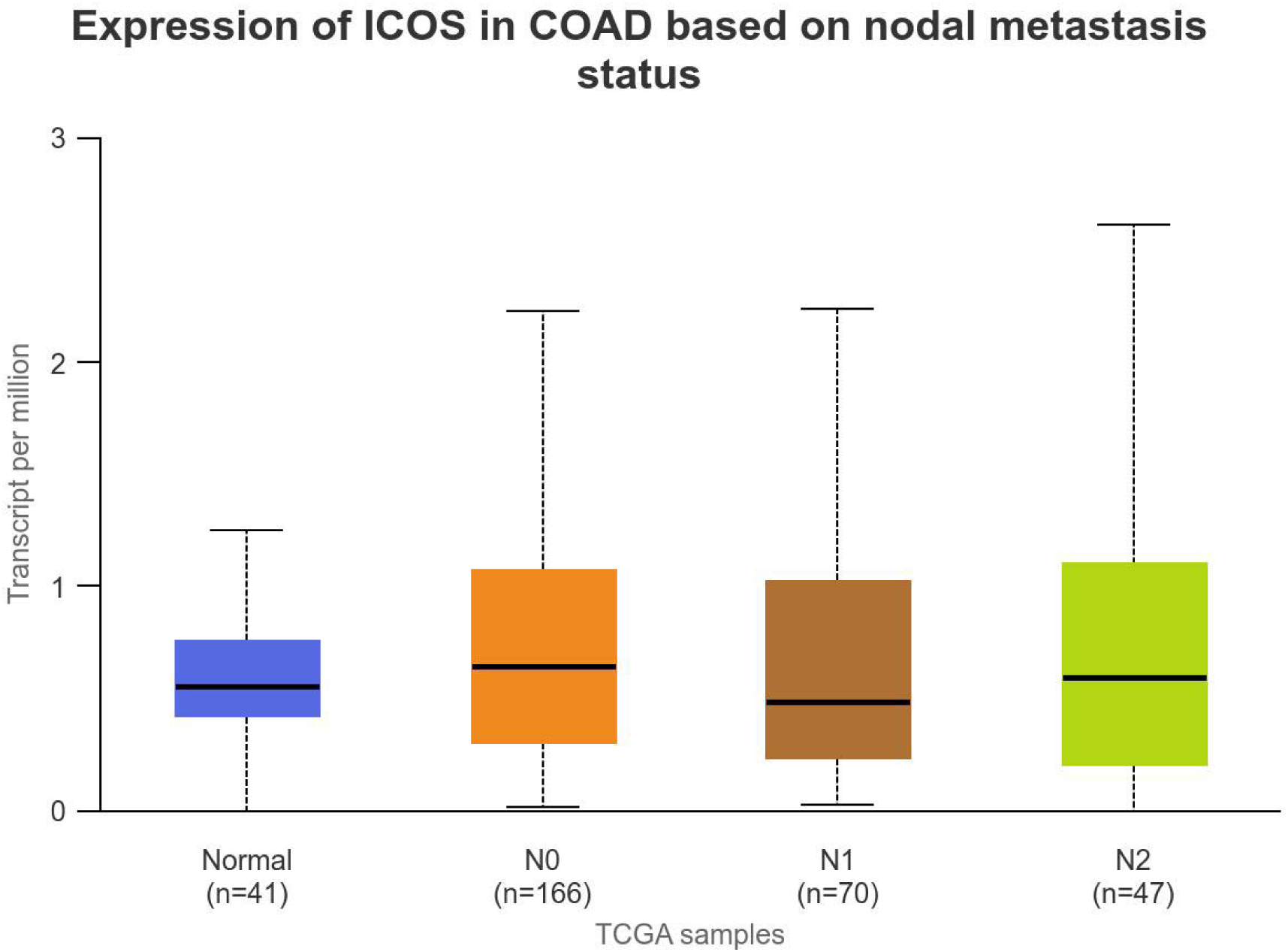
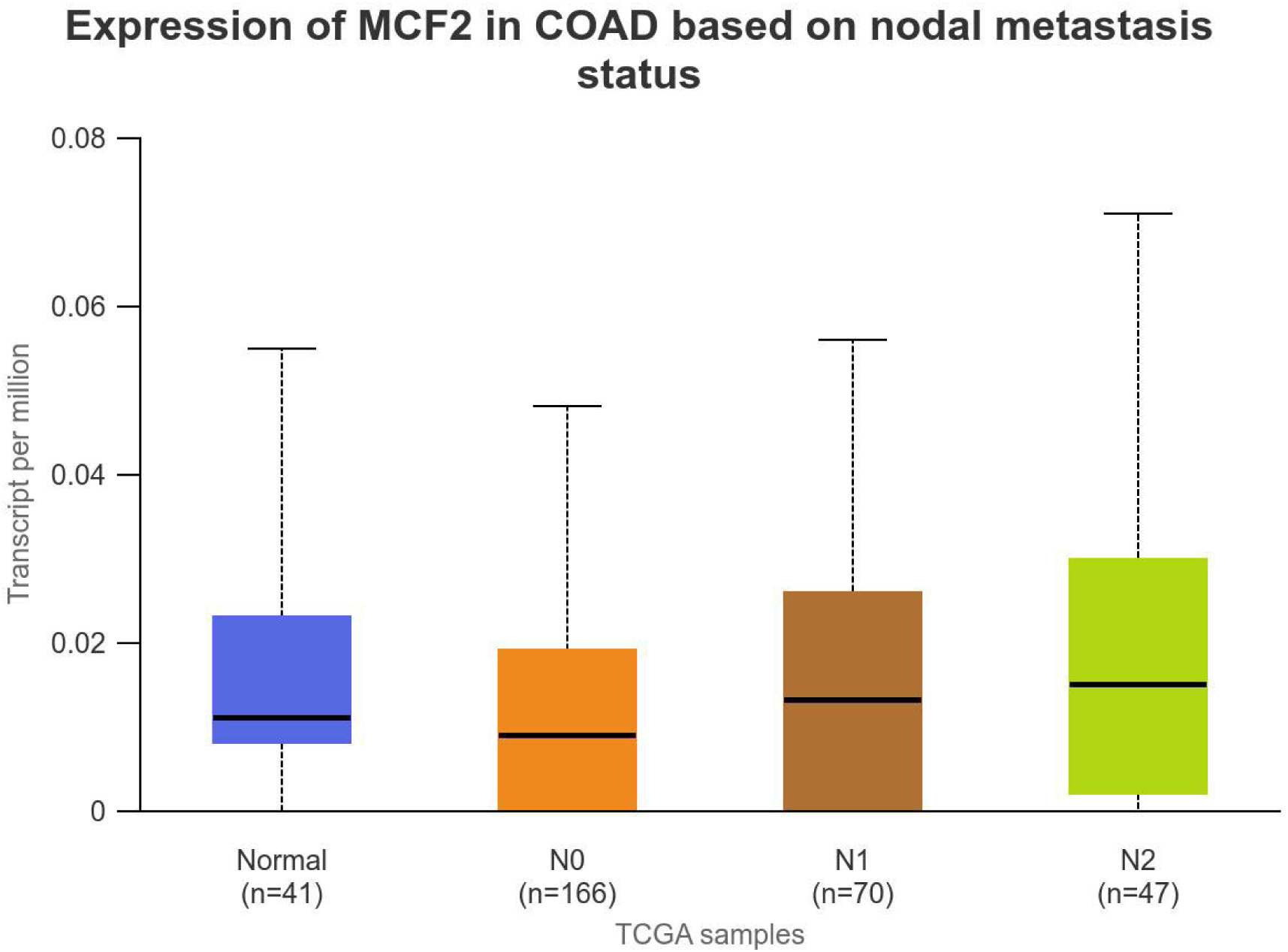
Prognostic characteristics of hub genes in COAD datasets. (a) The mRNA expression levels and overall survival rates of hub genes—CXCL8, FOXC2, ICOS, and MCF2—were analyzed in colon adenocarcinoma (COAD) patients using data from The Cancer Genome Atlas (TCGA) via the GEPIA database. Expression levels were compared between COAD patient samples (orange) and normal samples (blue), with statistical significance assessed through a Student’s t-test. Results indicated that these hub genes were significantly overexpressed in COAD patients compared to normal samples (*p < .001). b) Survival plots for CXCL8, FOXC2, ICOS, and MCF2 were generated to evaluate the relationship between hub gene expression levels and overall survival in COAD patients. The plots display two groups: high expression levels (orange line) and low expression levels (blue line). The horizontal axis represents time in months, while the vertical axis shows the survival rate percentage. Survival probabilities were estimated using the Kaplan-Meier method, with statistical significance assessed by the log-rank test (a) mRNA expression levels for different stages (I-IV) of CXCL8, FOXC1, ICOS, and MCF2 were obtained from the GEPIA database. b) The nodal metastatic status (N0–N3) for hub genes FOXC1, ICOS, and MCF2 was obtained from the UALCAN database and compared with normal CRC patient samples. Differences were considered statistically significant at a level of p < .001.

### Identification of mutation in hub genes

To investigate the genetic alterations of hub genes in colorectal cancer, we utilized the cBioPortal database. The expression levels and mutation status of the hub genes CXCL8, FOXC1, ICOS, and MCF2 were analyzed across 20 colorectal cancer studies, which included a total of 5,855 samples. Figure 7(a) illustrates the frequencies and types of genetic mutations in these hub genes within colorectal cancer samples. The analysis revealed that mutations were present in approximately 4% of the 20 cases examined. The mutation plots for the hub genes CXCL8, FOXC1, ICOS, and MCF2 in colorectal cancer indicate a mutation rate exceeding 1.0% in patient samples obtained from The Cancer Genome Atlas (TCGA) datasets. Additionally, the expression levels of the four hub genes (CXCL8, FOXC1, ICOS, and MCF2) in colorectal cancer were validated through immunostaining data obtained from the Human Protein Atlas (HPA) database, as illustrated in Figure 7(b). The immunostaining results reinforce the conclusion that CXCL8, and ICOS are significantly upregulated in colorectal cancer tissues. These findings contribute to the growing body of evidence that indicates genetic alterations in these hub genes are associated with the clinical characteristics of patient samples, suggesting their potential involvement in the initiation and progression of colorectal carcinogenesis.

**Figure 7.**
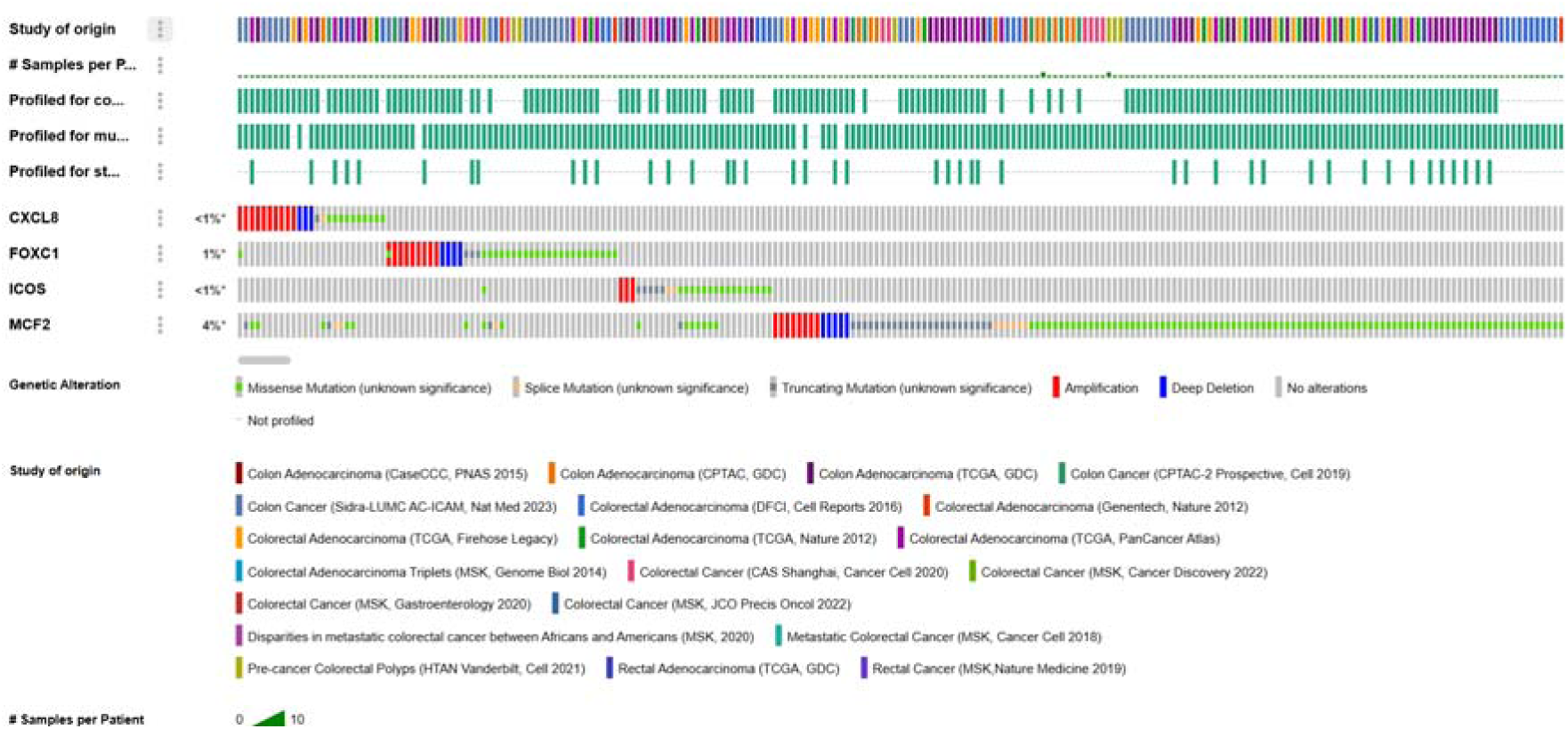
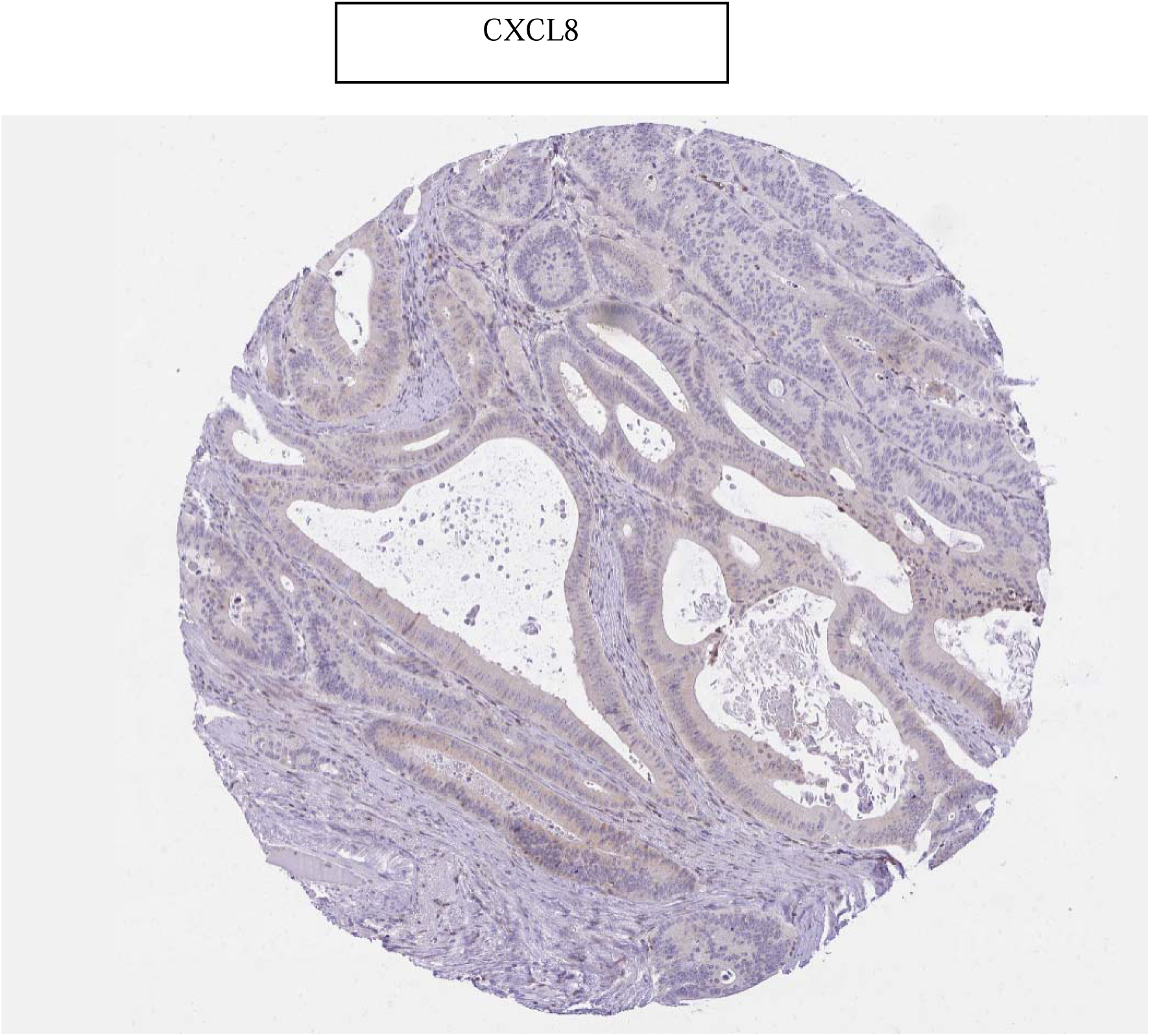
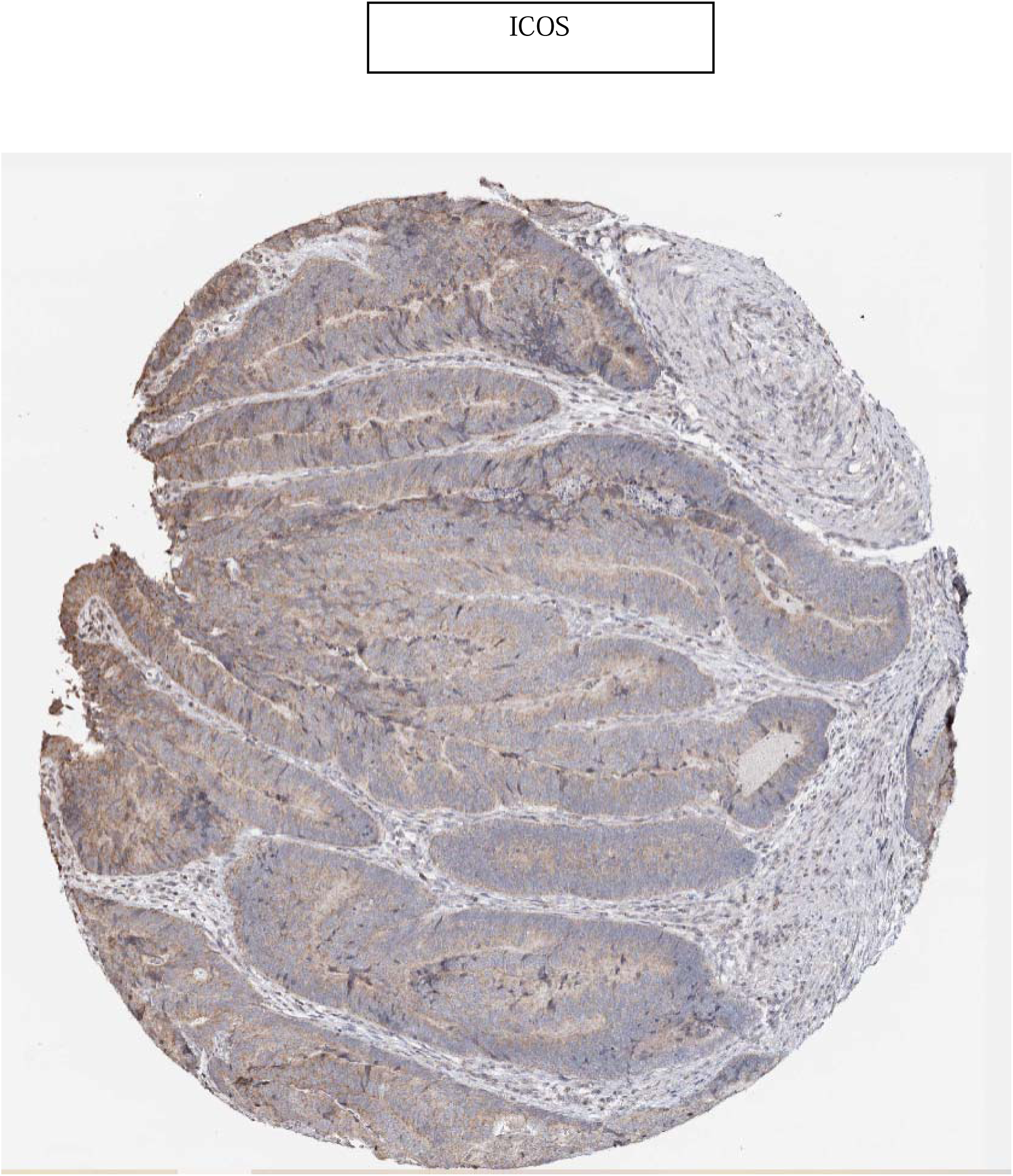
Alteration and validation of hub genes a) Gene mutation frequencies of hub genes CXCL8, FOXC1, ICOS, and MCF2 in colon adenocarcinoma (COAD) were retrieved from TCGA datasets via the cBioPortal database. Mutation types are represented as follows: red bars for gene amplifications, blue bars for deep deletions, green bars for missense mutations, and gray bars for truncating mutations. (b) Immunohistochemistry images of CXCL8, and ICOS in tumor tissues were obtained from the Human Protein Atlas database.

## Discussion

Colorectal cancer (CRC) is the second leading cause of cancer-related death in the United States, responsible for over 50,000 reported deaths in 2024, following lung and bronchus cancer. It is estimated that 150,000 new cases will be diagnosed annually, with a slight predominance among males compared to females (31) Microarray technology is considered a powerful tool for analyzing gene expression in colorectal cancer, providing insights into carcinogenesis, diagnosis, and treatment (32,33). It also unravels the complex biology of CRC, contributing to improved patient care through personalized medicine (34).

Analysis of the GSE164191 dataset identified 191 genes with altered expression levels in colorectal cancer (CRC) tissues compared to normal tissues, with 89 genes upregulated and 56 downregulated. This significant differential expression, defined by a p-value < 0.05 and |logFC| > 1, highlights the complexity of gene regulation in CRC. Identifying these differentially expressed genes (DEGs) is essential, as they hold potential as diagnostic biomarkers and therapeutic targets.

Key upregulated genes, including CXCL8, FOXC1, and MMP9, are known to play roles in tumor progression and metastasis. On the other hand, downregulated genes may represent loss of function in pathways crucial for cellular stability, possibly contributing to tumorigenesis. This identification of DEGs underscores the need to explore their specific roles in CRC.

Functional enrichment analysis using the DAVID tool offered insight into biological processes, cellular components, and molecular functions associated with these DEGs. Upregulated genes were primarily involved in transcription regulation, immune response, and cell adhesion—processes likely disrupted in CRC. For example, the involvement of DEGs in RNA polymerase II-mediated transcription suggests that abnormal gene expression may drive oncogenesis.

Downregulated genes, in contrast, were linked to essential functions like calcium-dependent cell-cell adhesion and glucose homeostasis, both critical for tissue integrity and metabolic stability. The disruption of these processes may promote tumor invasion and metastasis, emphasizing the functional importance of these DEGs.

Constructing a protein-protein interaction (PPI) network for the DEGs showed strong connections among them, forming a network with 131 nodes and 52 edges. Hub genes such as CXCL8, FOXC1, and MMP9, which were highly connected within this network, play pivotal roles in CRC biology. This analysis of the PPI network underlines the interconnected nature of these genes, pointing to possible pathways for therapeutic intervention.

Tools like STRING and FunRich were used to analyze protein-protein interactions, offering a broader view of the molecular landscape in CRC. This approach helps pinpoint regulatory nodes that may influence tumor behavior, which is key to developing therapies that disrupt these networks and halt tumor growth.

Survival analysis through databases like KM Plotter and GEPIA found that high expression levels of certain hub genes, especially CXCL8, FOXC1, ICOS, and MCF2, were linked to lower survival rates in CRC patients. These findings suggest these genes could serve as prognostic markers, associated not only with tumor development but also with disease progression.

Validation across multiple databases strengthened the case for these genes as significant prognostic indicators. Additionally, the analysis showed that these hub genes have mutation rates above 1% in patient samples, implying that genetic changes may enhance their oncogenic effects. This highlights the importance of integrating both genetic and expression data to fully understand the roles of these genes in CRC.

## Conclusion

The findings from this study provide a comprehensive overview of the dysregulated genes in colorectal cancer and their potential roles in tumor biology. The identification of DEGs, coupled with functional enrichment and PPI network analyses, offers valuable insights into the molecular mechanisms underlying CRC. Furthermore, the prognostic significance of hub genes emphasizes the need for further research to explore their potential as therapeutic targets and biomarkers for CRC. Future studies should focus on elucidating the specific roles of these genes in CRC progression and their interactions within the broader context of tumor biology.

